# Intraflagellar transport complex B proteins regulate the Hippo effector Yap1 during cardiogenesis

**DOI:** 10.1101/777128

**Authors:** Marina Peralta, Katerina Jerabkova, Tommaso Lucchesi, Laia Ortiz Lopez, Benjamin Vitre, Dong Han, Laurent Guillemot, Chaitanya Dingare, Izabela Sumara, Nadia Mercader, Virginie Lecaudey, Benedicte Delaval, Sigolène M. Meilhac, Julien Vermot

**Affiliations:** Institut de Génétique et de Biologie Moléculaire et Cellulaire, Illkirch, France; Centre National de la Recherche Scientifique, UMR7104, Illkirch, France; Institut National de la Santé et de la Recherche Médicale, U964, Illkirch, France; Université de Strasbourg, Illkirch, France; Imagine - Institut Pasteur, Laboratory of Heart Morphogenesis, Paris, France; INSERM UMR1163, Université Paris Descartes, Paris, France; Sorbonne Université, Collège Doctoral, F-75005, Paris, France; Centre de Recherche en Biologie cellulaire de Montpellier, (CRBM), CNRS, Univ. Montpellier, Montpellier, France; Institute for Cell Biology and Neurosciences, Goethe University of Frankfurt, Germany; Institute of Anatomy, University of Bern, Switzerland; Centro Nacional de Investigaciones Cardiovasculares CNIC, Madrid, Spain

## Abstract

Cilia and the intraflagellar transport (IFT) proteins involved in ciliogenesis are associated with congenital heart diseases (CHD). However, the molecular links between cilia, IFT proteins and cardiogenesis are yet to be established. Using a combination of biochemistry, genetics, and live imaging methods, we show that IFT complex B proteins (Ift88, Ift54 and Ift20) modulate the Hippo pathway effector YAP1 in zebrafish and mouse. We demonstrate that this interaction is key to restrict the formation of the proepicardium and the myocardium. *In cellulo* experiments suggest that IFT88 and IFT20 interact with YAP1 in the cytoplasm and functionally modulates its activity, identifying a molecular link between cilia related proteins and the Hippo pathway. Taken together, our results highlight a novel role for IFT complex B proteins during cardiogenesis and shed light on an unexpected mechanism of action for ciliary proteins in YAP1 regulation. These findings provide mechanistic insights into a non-canonical role for cilia related proteins during cardiogenesis.

## Introduction

Primary cilia are immotile microtubule-based organelles, well known for being both chemical and/or mechanical sensors (1). Disruption of cilia function causes multiple human syndromes known as ciliopathies (2). Cilia are required for cardiac development and mutations in cilia-related proteins have been linked to congenital heart diseases (CHDs) (3–6). Nevertheless, the specific role of cilia and cilia-related proteins during cardiogenesis is still unclear.

Intraflagellar transport (IFT) proteins are required for the transport of cilia components along the axoneme and are thus essential for cilia formation and function (7). The role of IFT proteins during ciliogenesis is highly conserved across organisms (8). The IFT machinery is composed of two biochemically distinct subcomplexes, IFT-A and IFT-B (8). The IFT complex B member IFT20, which localizes inside the cilium and at the Golgi complex in mammalian cells, participates in the sorting and/or transport of membrane proteins for the cilia (9). Within the IFT complex B, IFT20 interacts with IFT54/Elipsa (10, 11), and IFT88 which is essential for flagellar assembly in *Chlamydomonas* and ciliogenesis in vertebrates (12). Mutations in IFT proteins lead to ciliary defects and to several ciliopathies (2). Importantly, some IFT proteins have been identified in CHDs(4). IFT proteins also display non-canonical, cilia-independent functions (13, 14). IFT88, for example, is needed for spindle orientation or cleavage furrow ingression in dividing cells (15, 16) and regulates G1-S phase transition in non-ciliated cells (17). IFT20, together with IFT88 and IFT54, plays a role in the establishment of the immune synapse in T-lymphocytes lacking cilia (18, 19). These observations suggest that IFT proteins could play a key role in embryonic cardiogenesis through both cilia-dependent and – independent functions.

The Hippo signaling mediators, YAP1/WWTR1 (TAZ), constitute a key signaling pathway for the regulation of cardiac development (20–23) and cardiac regeneration (24, 25) in vertebrates. For example, YAP1/WWTR1 (TAZ) are required for epicardium and coronary vasculature development (26) in mice. Changes in cell shape, substrate stiffness and tension forces activate a phosphorylation-independent YAP1/WWTR1 (TAZ) modulation (27) which is mediated by the Motin family (AMOT, AMOTL1, and AMOTL2) (28, 29). Motin proteins bind to YAP/WWTR1 (TAZ) sequestering them in the cytoplasm in several cellular contexts (30–34). While it is known that ciliary proteins from the Nephrocystin family modulate the transcriptional activity of YAP1/WWTR1 (TAZ) (35–37) and that the Hippo kinases MST1/2-SAV1 pathway promotes ciliogenesis (38), the connection between the Hippo pathway and cilia function remains unclear.

Cardiogenesis involves an interplay between multiple tissue layers. The epicardium is the outermost layer covering the heart. This cardiac cell layer plays an essential role in myocardial maturation and coronary vessel formation during development (39–41) and has a crucial role during regeneration (42–44). Epicardial cells derive from the proepicardium (PE), an extra-cardiac transient cluster of heterogeneous cells (45). In mice, the PE is a single cell cluster located close to the venous pole of the heart (45), while in zebrafish is composed by two cell clusters, avcPE (the main source of cells) and vpPE (46). Proepicardial cells give rise to the epicardium, part of the coronary vasculature and intracardiac fibroblasts (45,47–49). The secreted signaling molecules of the bone morphogenetic protein (BMP) family are indispensable for PE formation (50–52). However, the regulation of PE development remains poorly understood.

Despite the increasing evidence for the role of IFT proteins in cell signaling, it has been difficult to pinpoint the exact function of cilia-related proteins outside the cilium. Without this key information, the question of whether cilia-related proteins can affect the developmental program independently of their cilia function remains unresolved. Here, we provide a combination of *in vivo* and *in vitro* analyses of IFT protein function showing that IFT complex B proteins can modulate the Hippo pathway effector Yap1. In particular, we show that IFT88 is required to restrict proepicardial and myocardial development through cytoplasmic activity.

## Results

### IFT complex B proteins regulate proepicardial development in zebrafish and mouse

Considering the importance of cilia during cardiogenesis, we assessed the role of several IFT complex B proteins during embryonic cardiogenesis *in vivo* focusing on the PE. In order to benefit from live imaging and genetics, we used zebrafish as a model organism. We performed live imaging focusing on the main PE cell source located near the atrioventricular canal (avcPE) in *ift88* (53) and *elipsa/ift54* (10) mutants (**Fig. 1 A-D**) (**Supp. 1 B, C**). Between 50 hpf and 57 hpf, we found that *ift88* mutants display multiple avcPE clusters (either 2 or 3), when controls have only one (**Movie 1**). Using a *wilms tumor 1 a* (*wt1a)* enhancer trap line that marks proepicardial and epicardial cells *(Et(−26.5Hsa.WT1-gata2:EGFP)^cn1^* hereafter *epi:GFP)* (46) we quantified avcPE and epicardial cell number and found that *ift88-/-* have increased avcPE cell numbers compared to their controls at 50 hpf. We additionally found that the size change is associated with an increased number of epicardial cells number in *ift88-/-* embryos (**Supp. 2 A**). Similarly, we found that *elipsa-/-* also showed bigger avcPE compared to controls (**Fig. 1 D**) (**Supp. 1 A’’**). Taken together, these data indicate that *ift88* and *elipsa*/*ift54* are required to restrict PE size.

**Figure 1.**
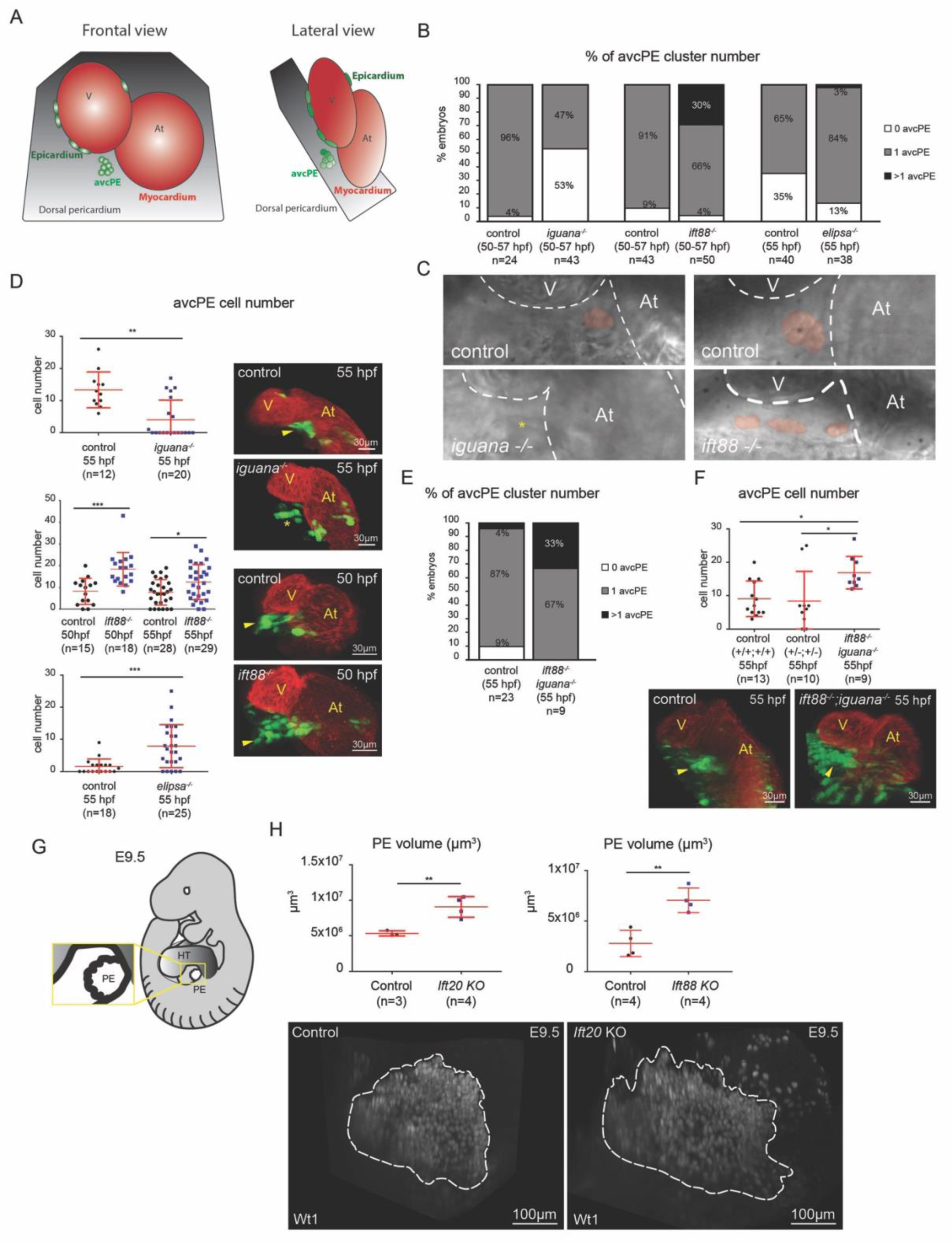
IFT complex B proteins regulate proepicardial development in zebrafish and mice. **(A)** Schematic representation of the zebrafish heart at 55 hpf in frontal and lateral views. The dorsal pericardium is shown in grey, the myocardium in red, the avcPE in light green and the epicardial cells in dark green. V, ventricle; At, atrium.**(B)** Percentage of avcPE cluster number in iguana (n=43), ift88 (n=50) and elipsa/ift54 (n=38) mutants and their controls (n=24; n=43; n=40 respectively), between 50 and 57 hpf. **(C)** High-speed avcPE imaging (digital red mask to improve visualization) of iguana and ift88 mutants and their control. **(D)** Graphs show avcPE cell number quantified on iguana (n=20), ift88 (n=18 and 29) and elipsa (n=25) mutants in epi:GFP background and their controls (n=12; n=15; n=28 n=18 respectively). (t-test iguana p-value 0.001; ift88 p-value 0.0002 and p-value 0.015 respectively; elipsa p-value 0.0005). 3D projections of whole mount immunofluorescence of hearts using an anti-myosin heavy chain antibody (red) and GFP (green) expression. Ventral views, anterior is to the top. Arrowheads mark avcPE and asterisk shows lack of avcPE. **(E)** Percentage of avcPE cluster number on iguana-/-, ift88-/-, epi:GFP (n=9) and controls (n=23) at 55 hpf. **(F)** Graph shows avcPE cell number quantified in wild type (n=13) and double heterozygous controls (n=10) (ift88+/+; iguana+/+; epi:GFP and ift88^+/-^; iguana^+/-^; epi:GFP) and double ift88; iguana mutants (n=9) (ift88-/-; iguana-/-; epi:GFP) (Kruskal-Wallis p-value 0.014). 3D projections of whole mount immunofluorescence of hearts using myosin heavy chain antibody (red) and GFP (green) expression. Ventral views, anterior is to the top. Arrowheads mark avcPE. V, ventricle; At, atrium. **(G)** Schematic representation of an E9.5 mouse embryo. Heart tube (HT) in dark grey and PE in white. Section of the PE represented inside the yellow box. **(H)** Left side graph shows quantification of PE volume (µm3) in Ift20 KO (n=4) and control mice (n=3) (control 5.35×10^6^µm^3^±3.63×10^5^; Ift20 KO 9.06×10^6^µm^3^±1.46×10^6^) (t-test p-value 0.008) and that of Ift88 KO (n=4) and control embryo (n=4) on the right (control 2.78×10^6^µm^3^±1. 3×10^6^; Ift88 KO 7.05×10^6^µm^3^±1.2×10^6^) (t-test p-value 0.003). In the lower panel, 3D projections of immunofluorescence whole-mount performed on control and Ift20 KO embryos. PE marked using anti-Wt1 antibody. White dotted shapes enclose the PE area. In all graphs, red bars indicate mean ± standard deviation.

To determine whether primary cilia are required for PE formation, we took advantage of the *iguana/dzip* (54) mutant, a well-established cilia mutant lacking primary cilia (55). We confirmed the absence of primary cilia in the pericardial cavity of the *iguana-/-* using the cilia reporter *actb2:Mmu.Arl13b-GFP* (56) (**Supp. 2 C-E**) (**Movie 2**). Zebrafish *ift* genes are expressed maternally (57) (**Supp. 2 E**) and complete removal of both zygotic and maternal expression of *ift88* leads to the ablation of primary cilia resembling the *iguana* mutant phenotype (58). Live imaging revealed that a significant fraction of *iguana* mutants lacked an avcPE between 50 hpf and 57 hpf (**Fig. 1 B**) (**Movie 3**). At 55 hpf, *iguana-/-* displayed decreased avcPE cell numbers (**Fig. 1 D**) (**Supp. 1 A and 2 A’**). Since the *iguana* mutants presented a phenotype opposite to that of the *ift88* mutants, we analyzed the *ift88-/-; iguana-/-; epi:GFP* double mutant to elucidate whether the differences could be due to a new cilia-independent function. The *ift88-/-; iguana-/-;* double mutant shows multiple avcPE cluster formation and an increased avcPE cell number, reminiscent of *ift88* loss of function (**Fig. 1 E, F**) (**Supp. 1 D**). We conclude that Ift88 modulates the PE cell number independently of its cilia function. We next assessed whether the role of IFT complex B proteins during PE formation was conserved in mammals. We analyzed the PE in *Ift20* and *Ift88* knock-out (KO) mice at mouse embryonic day (E) 9.5 (**Fig. 1 G, H**) (**Supp. 3 A, B**). First, we performed immunofluorescence using an anti-Arl13b antibody to quantify the percentage of ciliated PE cells in *Ift20*, *Ift88 KO* and *wild type* mice (**Supp. 3 C-C’’**). We found that, while over 60% of the *wild type* PE cells were ciliated, in *Ift88 KO* mice only 3% of PE cells were ciliated and *Ift20 KO* embryos lack cilia (**Supp. 3 C’’**). We compared the PE volume of *Ift20 KO* mice to their controls, using an anti-Wt1 antibody (a well-established PE marker (59)). Mutant embryos showed increased PE volume compare to controls (**Movie 5, 6**). Likewise, the PE volume in *Ift88 KO* was also increased compared to controls (**Fig. 1 H**).Together, these results suggest that IFT protein function is conserved during PE formation in mouse and zebrafish and that, in zebrafish, IFT88 contributes to this process independently of its ciliary function.

To test whether IFT complex B affected other cardiac tissues, we focused in the myocardium. Cell quantification showed increased myocardial cell number in *ift88-/-* compared to their controls at 50 hpf (**Fig. 2 A, A’**) and 55 hpf (**Supp. 4 A**). Interestingly, the increment was mostly affecting the atrium. Similar data were obtained with the *elipsa* mutants at 55hpf (**Fig. 2 B**). These results led us to conclude that the absence of IFT complex B proteins also affects the myocardial cells.

**Figure 2.**
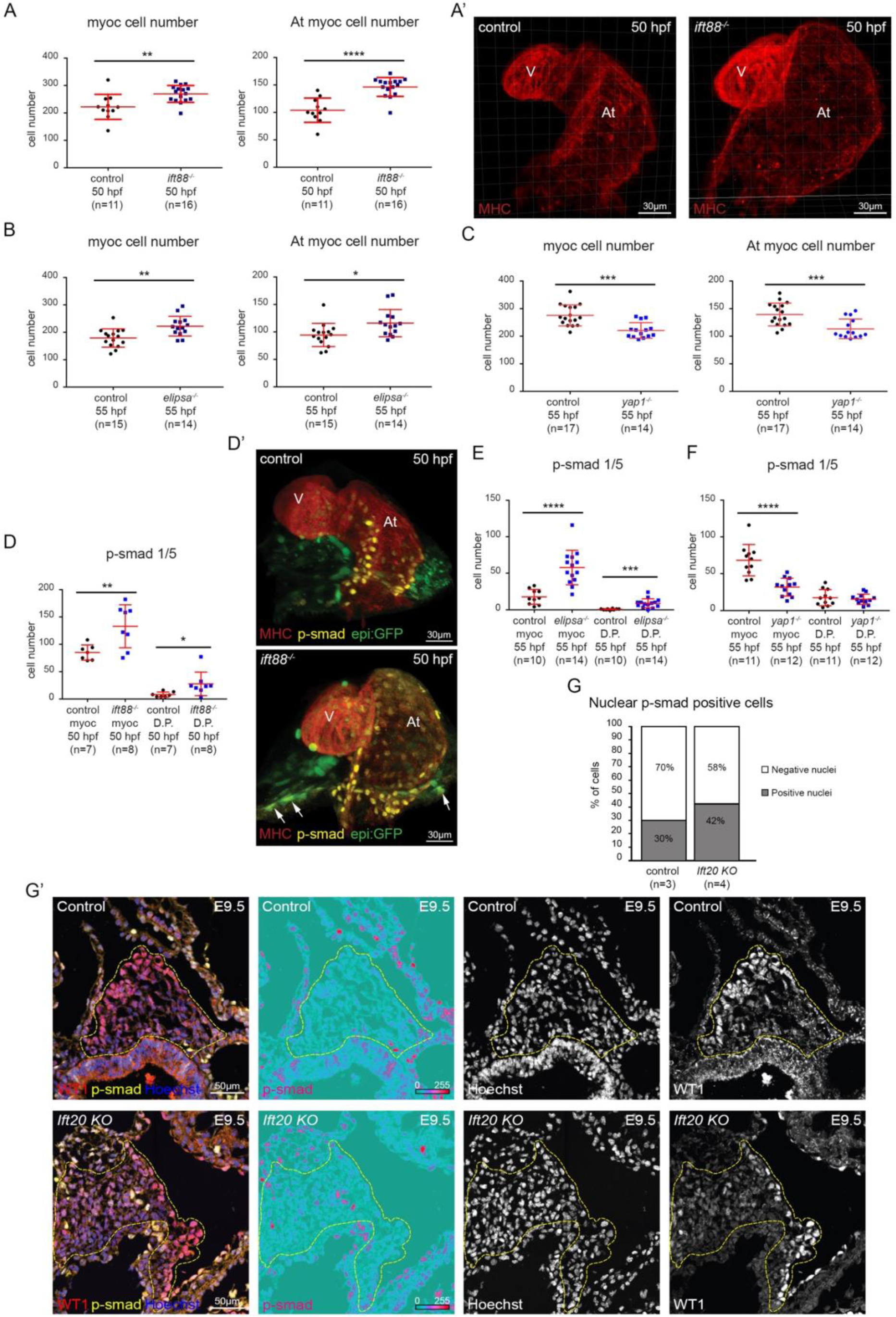
BMP signaling is increased in ift88, elipsa/ift54 and Ift20 mutants. **(A)** Graphs show total myocardial and atrial-myocardial cell number quantified in ift88 (n=16) mutants and controls (n=11) at 50 hpf. (t-test total myocardium p-value 0.0034; atrial myocardium p-value <0.0001). **(A’)** 3D projections of whole mount immunofluorescence of hearts using myosin heavy chain antibody (MHC) (red). Ventral views, anterior is to the top. **(B)** Graphs show total myocardial and atrial-myocardial cell number quantified in elipsa (n=14) mutants and controls (n=15) at 55 hpf. (t-test total myocardium p-value 0.0026; atrial myocardium p-value 0.017). **(C)** Graphs show total myocardial and atrial-myocardial cell number quantified in yap1 (n=14) mutants and controls (n=17) at 55 hpf. (t-test total myocardium p-value 0.0001; atrial myocardium p-value 0.0009). **(D)** Graph shows number of p-smad1/5 positive cells in the myocardium and dorsal pericardium (DP) quantified in ift88 mutants (n=8) and controls (n=7) at 50 hpf. Myocardium (t-test p-value 0.009) and DP (t-test p-value 0.036). **(D’)** 3D projections of whole mount immunofluorescence of hearts using myosin heavy chain antibody (MHC) (red), epi:GFP (green) and p-smad1/5 (yellow) antibody. Arrows mark dorsal pericardial cells positive for epi:GFP and p-smad1/5. Ventral views, anterior is to the top. **(E)** Graph shows number of p-smad1/5 positive cells in the myocardium and dorsal pericardium (DP) quantified in elipsa mutants (n=14) and controls (n=10) at 55 hpf. Myocardium (t-test p-value < 0.0001) and DP (t-test p-value 0.0004). **(F)** Graph shows number of p-smad1/5 positive cells in the myocardium and dorsal pericardium (DP) quantified in yap1 mutants (n=12) and controls (n=11) at 55 hpf. Myocardium (t-test p-value < 0.0001) and DP (t-test p-value 0.599). **(G)** Graph shows that the percentage of p-smad 1/5/9 positive PE cells is higher in Ift20 KO (n=4 embryos: 1862 nuclei analyzed) compared to control (n=3 embryos: 1414 nuclei analyzed) mice (Chi-square test of homogeneity =51.593, p-value 6.829e-13 on 1 degree of freedom). **(G’)** Control and Ift20 KO immunofluorescence confocal cryosections labelled with WT1 (red), p-smad 1/5/9 (yellow) and Hoechst (blue) at E9.5. Yellow dotted lines enclose the PE area. Individual channels are displayed for p-smad 1/5/9 (signal is shown as ice LUT to facilitate the visualization of signal intensity, where green is the minimum and red is the maximum), Hoechst (white) and WT1 (white). In all graphs, red bars indicate mean ± standard deviation. V, ventricle; At, atrium. p-smad, p-smad1/5 in panel D**’**and p-smad1/5/9 in panel G**’**.

### BMP signaling is increased in ift88, elipsa/ift54 and Ift20 mutants during PE development

As it is known in zebrafish that overexpressing BMP increases PE size (50), we investigated whether *ift88* loss influences BMP signaling. We first assessed *bmp4* expression by *in situ* hybridization in *ift88* and *elipsa* mutants. We found that *ift88* mutant embryos display ectopic expression of *bmp4* in the dorsal pericardium (DP) at 48 hpf (**Supp. 4 B**). To validate that the upregulation is functional *in vivo*, we quantified cellular BMP activity by using p-smad 1/5 as a readout and confirmed that the absence of Ift88 leads to increased BMP signaling activity (**Fig. 2 D, D’**). Additionally, more myocardial cells were positive for p-smad 1/5 in *ift88-/-* than in controls, especially in the venous pole (**Fig. 2 D, D’**) (**Supp. 4 C**). By 55 hpf, *bmp4* expression at the atrioventricular canal myocardium is reduced and the expression at the venous pole is almost undetectable in controls. By contrast, *ift88* mutants presented strong *bmp4* expression in both domains (**Supp. 4 B**). Myocardial p-smad 1/5 was also increased in *ift88-/-* compared to controls, but not in the DP (**Supp. 4 C, D**). We reached similar conclusions when analyzing the *elipsa* mutants at 55 hpf (**Fig. 2 E**) (**Supp. 4 B-C**). To confirm that the regulation of BMP signaling by IFT is conserved in vertebrates, we next analyzed p-smad 1/5/9 in *Ift20* and *Ift88 KO* and control mice at E9.5 and found that only the *Ift20* mutants displayed a higher percentage of p-smad-positive cells compared to controls (**Fig. 2 G, G’**). (**Supp. 3 D**). Together, these results suggest that Ift88 and Ift54 modulate BMP signaling activity in the myocardium and the pericardium of the zebrafish, and that IFT20 plays a similar role in the mouse PE.

### Yap1-Tead activity is increased in ift88, elipsa/ift54 and Ift20 mutants during PE development

YAP1/WWTR1, which are important regulators of tissue size (60), are essential for epicardium and coronary vasculature development (26). In addition, Yap1 participates in the regulation of Bmp signaling during secondary heart field development (21). This led us to assess if Yap1/Wwtr1-Tead is active during PE formation. We first analyzed Yap1/Wwtr1-Tead activity in PE cells using the *4xGTIIC:d2GFP* reporter line (61). We found that the reporter was active in the PE cluster, myocardium, pericardium and epicardial cells at 55 hpf (**Supp. 5 A**). Time-lapse analysis allowed us to evaluate the dynamics of Yap1/Wwtr1-Tead activity during PE development between 51 and 60 hpf (**Movie 4**) (**Supp. 5 B**). GFP quantification in PE and pericardial cells showed higher Yap1/Wwtr1-Tead activity (GFP average intensity) in the PE cells than in the pericardial cells (75% of the embryos) (**Supp. 5 B**) (**Supp. Table 1**). Thus, Yap1/Wwtr1-Tead is active in PE cells during PE development.

We next tested whether the increased avcPE cell number was due to abnormal Yap1 activity, we performed immunofluorescence to quantify the number of nuclear Yap1-positive cells in *ift88* and *elipsa* mutants (**Fig. 3A, A’**) (**Supp. 6 A-B’**). At 55 hpf, *ift88-/-* showed more nuclear Yap1-positive cardiomyocytes than controls (**Fig. 3 A**), especially in the atrium (**Supp. 6 A**). Similar data were obtained with the *elipsa* mutants (**Fig. 3 A**) (**Supp. 6 A**). To further validate the link between increased PE size and Yap1-Tead activity, we treated the *ift88* mutants with the drug Verteporfin, which binds to YAP and changes its conformation, blocking its interaction with TEAD (62). We first performed time-lapse imaging on Verteporfin treated *4xGTIIC:d2GFP* embryos at 55 hpf to assess Verteporfin specificity. We measured the Yap/Wwtr1-Tead activity (GFP average intensity) in the same PE and pericardial cells at several timepoints: before adding Verteporfin (t0), after 2 hours treatment with Verteporfin (5µM), and 45 min after washing out (**Supp. 5 C**). We found that Yap/Wwtr1-Tead activity is significantly decreased with Verteporfin, confirming the specificity of the drug on the embryo. We next treated *ift88-/-* and control (*ift88+/+* and *^+/-^)* embryos with Verteporfin (5µM) from 50 hpf to 55 hpf (**Fig. 3 B**). Control embryos treated with Verteporfin showed smaller avcPE compared to untreated controls. Verteporfin-treated *ift88-/-* embryos presented fewer avcPE cell numbers than non-treated *ift88-/-* embryos. Interestingly, non-treated controls and treated *ift88* mutants did not show significant differences in avcPE cell number. Thus, the increase in avcPE cell number induced in the absence of Ift88 was rescued by Verteporfin treatment. These results suggest that Ift88 requires Yap1 activity to modulate the PE size restriction.

**Figure 3.**
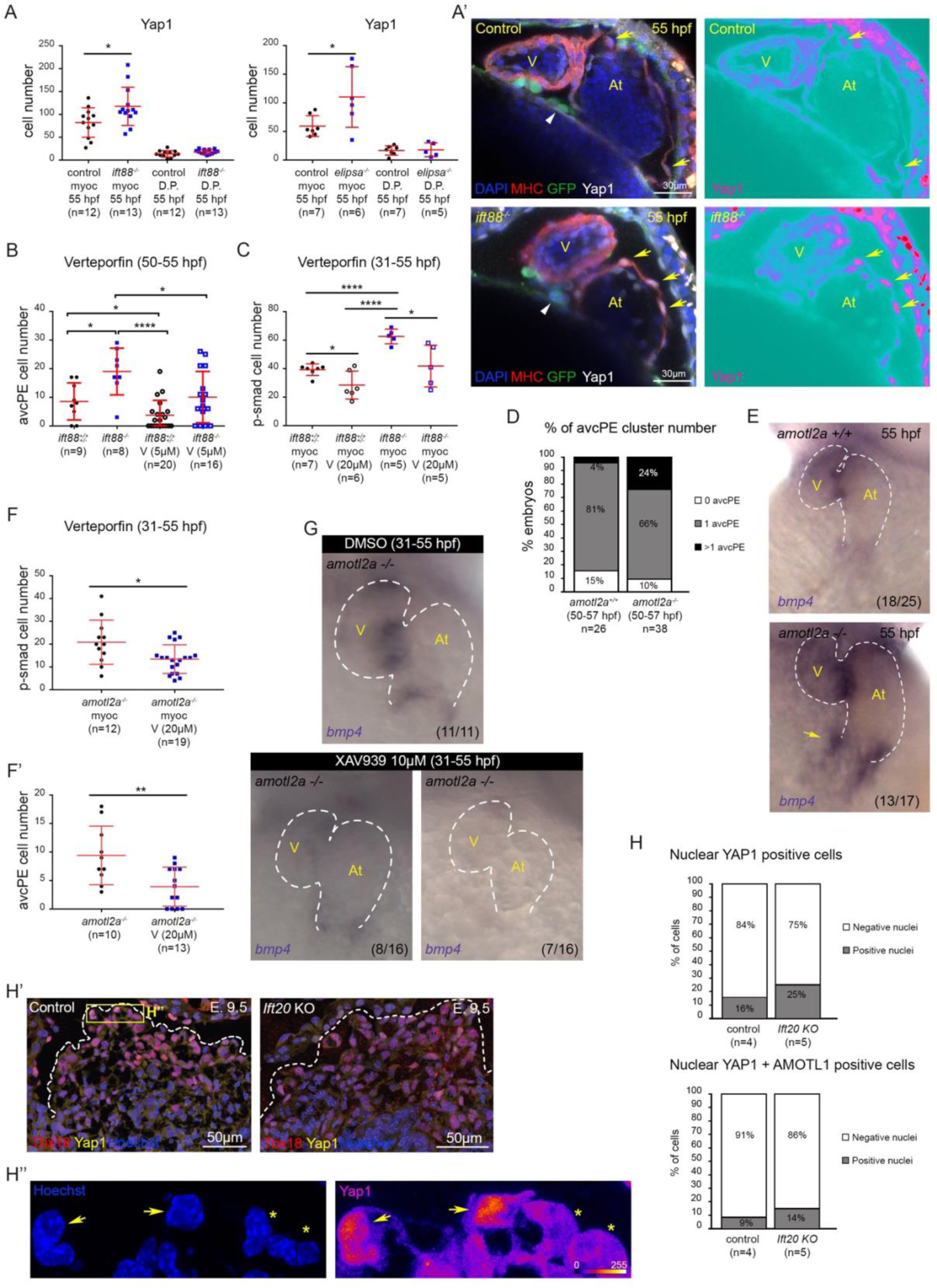
Yap1-Tead activity is increased in ift88, elipsa/ift54 and Ift20 mutants during proepicardium development. **(A)** Graphs show number of Yap1-positive cells in the myocardium and dorsal pericardium (DP) quantified in ift88 (n=13), elipsa (n=6) and their controls (n=12 and n=7 respectively). ift88 mutants show increased Yap1-positive myocardial cell numbers (t-test p value 0.03) and a tendency towards higher Yap1 positive DP cell numbers (t-test p value 0.07). elipsa mutants show higher Yap1-positive myocardial cell numbers (t-test p value 0.036). **(A’)** Control and ift88-/-, epi:GFP immunofluorescence confocal sections labelled with anti-myosin heavy chain antibody (MHC) (red), GFP (green),-Yap1 (white) and DAPI (blue) at 55 hpf. Ventral view, anterior is to the top. Individual channel is displayed for Yap1 (signal is shown as ice LUT to facilitate the visualization of signal intensity, where green is the minimum and red is the maximum). Yellow arrows mark nuclear Yap1-positive atrial myocardial cells. White arrowheads mark the avcPE. **(B)** Graph shows avcPE cell number quantified in control (n=9), ift88-/-, epi:GFP (n=8) and Verteporfin (5µM)-treated ift88-/-, epi:GFP (n=16) and control (n=20) embryos (55hpf). Control embryos treated with Verteporfin showed smaller avcPE compared to untreated controls (t-test p-value 0.04). Verteporfin-treated ift88-/-; epi:GFP embryos presented lower avcPE cell numbers than non-treated ift88-/-; epi:GFP embryos (t-test p-value 0.027). ift88-/-; epi:GFP embryos showed bigger avcPE compared to untreated (t-test p-value 0.01) and treated controls (t-test p-value 0.0001). **(C)** Graph shows number of p-smad 1/5-positive cells in the myocardium quantified in control (n=7), ift88-/-, epi:GFP (n=5) and Verteporfin (20µM)-treated ift88-/-, epi:GFP (n=5) and control (n=6) embryos from 31 hpf to 55 hpf. Control embryos treated with Verteporfin showed decreased p-smad 1/5-positive cell numbers compared to untreated controls (t-test p-value 0.0219). Verteporfin-treated ift88-/-; epi:GFP embryos presented less p-smad 1/5-positive cells than non-treated ift88-/-; epi:GFP embryos (t-test p-value 0.0173). ift88-/-; epi:GFP embryos showed more p-smad 1/5-positive cells compared to untreated (t-test p-value <0.0001) and treated controls (t-test p-value <0.0001). **(D)** Percentage of avcPE cluster number in amotl2a+/+ (n=26) and amotl2a-/- (n=38) embryos between 50 and 57 hpf. **(E)** Whole mount bmp4 in situ hybridization in amotl2a+/+ (n=18/25) and amotl2a-/- (n=13/17) embryos (55hpf). Yellow arrow shows bmp4 overexpression. Ventral views, anterior is to the top. **(F)** Graph shows number of p-smad 1/5-positive cells in the myocardium quantified in Verteporfin (20µM)-treated amotl2a-/- (n=19) embryos from 31 to 55 hpf and untreated amotl2a-/- (n=12) embryos. Treated embryos showed decreased p-smad 1/5-positive cell numbers compared to untreated ones (t-test p-value 0.0148). **(F’)** Graph shows avcPE cell number quantified in Verteporfin (20µM)-treated amotl2a-/- (n=13) embryos from 31 to 55 hpf and untreated amotl2a-/- (n=10) embryos. Treated embryos showed decreased avcPE cell numbers compared to untreated ones (t-test p-value 0.0059). **(G)** Whole mount bmp4 in situ hybridization in XAV939 (10µM)-treated amotl2a-/- (n=16) from 31 to 55 hpf and untreated embryos (n=11). Treated embryos showed either decreased (n=8/16) or absent (n=7/16) bmp4 expression at the atrioventricular canal myocardium and the venous pole. Ventral views, anterior is to the top. **(H)** Graphs show percentages of YAP1-positive PE cells and double YAP1-AMOTL1-positive PE cells in Ift20 KO (n=5 embryos: 1196 nuclei analyzed) and control (n=4 embryos: 929 nuclei analyzed) mice at E9.5. The percentage of nuclear YAP1-positive PE cells (Chi-square test of homogeneity =25.354, p-value 4,77E-07 on 1 degree of freedom) and nuclear YAP1-AMOTL1-positive cells (Chi-square test of homogeneity =12,025, p-value 5,25E-04 on 1 degree of freedom) were higher in Ift20 KO than in control mice. **(H’)** Control and Ift20 KO immunofluorescence confocal cryosections labelled with TBX18 (red), YAP1 (yellow) and Hoechst (blue) at E9.5. White dotted lines enclose the PE area. **(H’’)** Zoomed region shows the difference between nuclear YAP1-positive cells (yellow arrows) and YAP1-negative cells (yellow asterisks). Hoechst signal (blue) highlights cell nuclei. YAP1 signal is shown as fire LUT to facilitate the visualization of signal intensity, where blue is the minimum and yellow is the maximum. In all graphs, red bars indicate mean ± standard deviation.

Additionally, to explore a possible role of Yap1 in the regulation of BMP signaling by IFT, we treated *ift88-/-* and control (*ift88+/+* and *^+/-^)* embryos with Verteporfin (20µM) from 31 hpf to 55 hpf (**Fig. 3 C**). We quantified cellular BMP activity by using p-smad 1/5 as a readout. Control embryos treated with Verteporfin showed decreased p-smad 1/5-positive myocardial cell number compared to untreated controls. Verteporfin-treated *ift88-/-* embryos presented fewer p-smad 1/5-positive myocardial cells than non-treated *ift88-/-* embryos. Remarkably, non-treated controls and treated *ift88* mutants did not show significant differences in p-smad 1/5-positive myocardial cells. Thus, the increase in BMP activity induced in the absence of Ift88 was rescued by Verteporfin treatment. Likewise, *in situ* hybridization performed in *elipsa-/-* and control (*elipsa+/+* and *^+/-^)* embryos treated with Verteporfin (20µM) from 31 hpf to 55 hpf displayed decreased *bmp4* expression (**Supp. 6 C**). Furthermore, treatment with the drug XAV939 (10µM), a tankyrase inhibitor that suppressed YAP-TEAD activity (63) also lead to a reduction of *bmp4* expression (**Supp. 6 C**). These results suggest that Ift88 and Ift54 require Yap1 activity to modulate BMP signaling activity in the myocardium of the zebrafish. To confirm the involvement of an increased Yap1 activity in the *ift88* mutant phenotype, we analyzed the avcPE in *angiomotin like 2a* (*amotl2a*) mutants (**Fig. 3 D, E**). In zebrafish, Amotl2a physically interacts with Yap1 leading to its cytoplasmic retention in a way that *amotl2a* mutants show upregulated Yap1 activity (nuclear Yap1) (33, 34). Accordingly, *amotl2a-/-* presented multiple avcPE formation. In addition, *bmp4* expression was increased in *amotl2a-/-* when compared to *amotl2a+/+* embryos at 55 hpf. These data are consistent with the *ift88* mutant phenotype and suggest that Ift88 modulates Yap1 activity to restrict PE size. To further validate the link between increased Yap1 and BMP signaling activity, we treated the *amotl2a* mutants with the drug Verteporfin (20µM) from 31 hpf to 55 hpf (**Fig. 3 F, F’**). *amotl2a-/-* embryos treated with Verteporfin showed decreased p-smad 1/5-positive myocardial cell number and avcPE cells compared to untreated *amotl2a-/-* embryos. Likewise, *bmp4* expression at the atrioventricular canal myocardium and at the venous pole were reduced in *amotl2a-/-* embryos treated with XAV939 (10µM) from 31 hpf to 55 hpf, when compared to untreated *amotl2a-/-* embryos (**Fig. 3 G**). Thus, Verteporfin and XAV939 treatments rescued the increase in BMP signaling induced in the absence of Amotl2a suggesting it acts through Yap1 activity. To further confirm the link between Yap1 activity and PE formation, we next analyzed the *yap1* mutants (33). At 55 hpf, *yap1-/-* showed decreased myocardial cell number compared to their controls (**Fig. 2 C**). Besides, fewer myocardial cells were positive for p-smad 1/5 in *yap1* mutants (**Fig. 2 F**). Surprisingly, avcPE cell number was similar between *yap1-/-* and their controls (**Supp. 2 B, B’**). Altogether, these results suggest that Ift88 and Ift54 modulate BMP signaling activity by tuning Yap1-Tead activity in the myocardium of the zebrafish. To assess if the upregulation of Yap1 activity observed in IFT zebrafish mutants was conserved in vertebrates, we analyzed YAP1 localization in the PE of *Ift20 KO* and control mice (**Fig. 3 H-H’’**). We quantified the proportion of nuclear YAP1-positive cells (i.e. cells with higher signal intensity inside the nucleus than in the cytoplasm) (**Fig. 3 H’’**). The percentage of nuclear YAP1-positive PE cells was higher in *Ift20 KO* than in control mice (**Fig. 3 H**). In mouse cardiomyocytes, YAP1 and AMOTL1 translocate to the nucleus together to modulate cell response (23). To assess AMOTL1 localization in PE cells, we performed AMOTL1 immunofluorescence. Consistent with the results obtained with an anti-YAP1 antibody, the percentage of nuclear AMOTL1-positive cells (i.e. cells with higher signal intensity inside the nucleus than in the cytoplasm) was increased in the *Ift20 KO* when compared to control mice (**Supp. 6 D**). We also found that the mutants showed an increase in the percentage of YAP1–AMOTL1 double positive cells (**Fig. 3 H**). Together, these data show that the increased PE size in *Ift20 KO* mice is associated with increased YAP1 activity.

### IFTs interact with YAP1 and regulate its activity

Taking advantage of cultured cells, we next explored the mechanism by which IFT proteins could regulate YAP1 activity. Immunofluorescence performed in HeLa cells transfected with IFT88-GFP, showed co-localization between IFT88 and YAP1 in the cytoplasm (**Supp. 7 A-C**). To confirm a potential physical interaction between IFT proteins with YAP1, we first performed Co-IP experiments using IFT88-GFP and YAP1-Myc in HeLa cells but never obtained any clear interaction. Considering that YAP1 often necessitates the scaffold protein Angiomotin-like 1 (Amotl1) (23), we next performed Co-IP experiment using HA-Amotl1, IFT88-GFP, and YAP1-Myc in HeLa cells. The experiments revealed a clear interaction between YAP1, Amotl1, and IFT88 (**Fig. 4 A**). Similarly, physical interaction between endogenous YAP1, Flag-Amotl1, and IFT20-GFP was observed in transfected HEK293 cells (**Fig. 4 A**). Together, these results reveal that IFT proteins, YAP1, and Amotl1 could function as part of a complex involved in the functional modulation of YAP1 activity *in vivo*.

**Figure 4.**
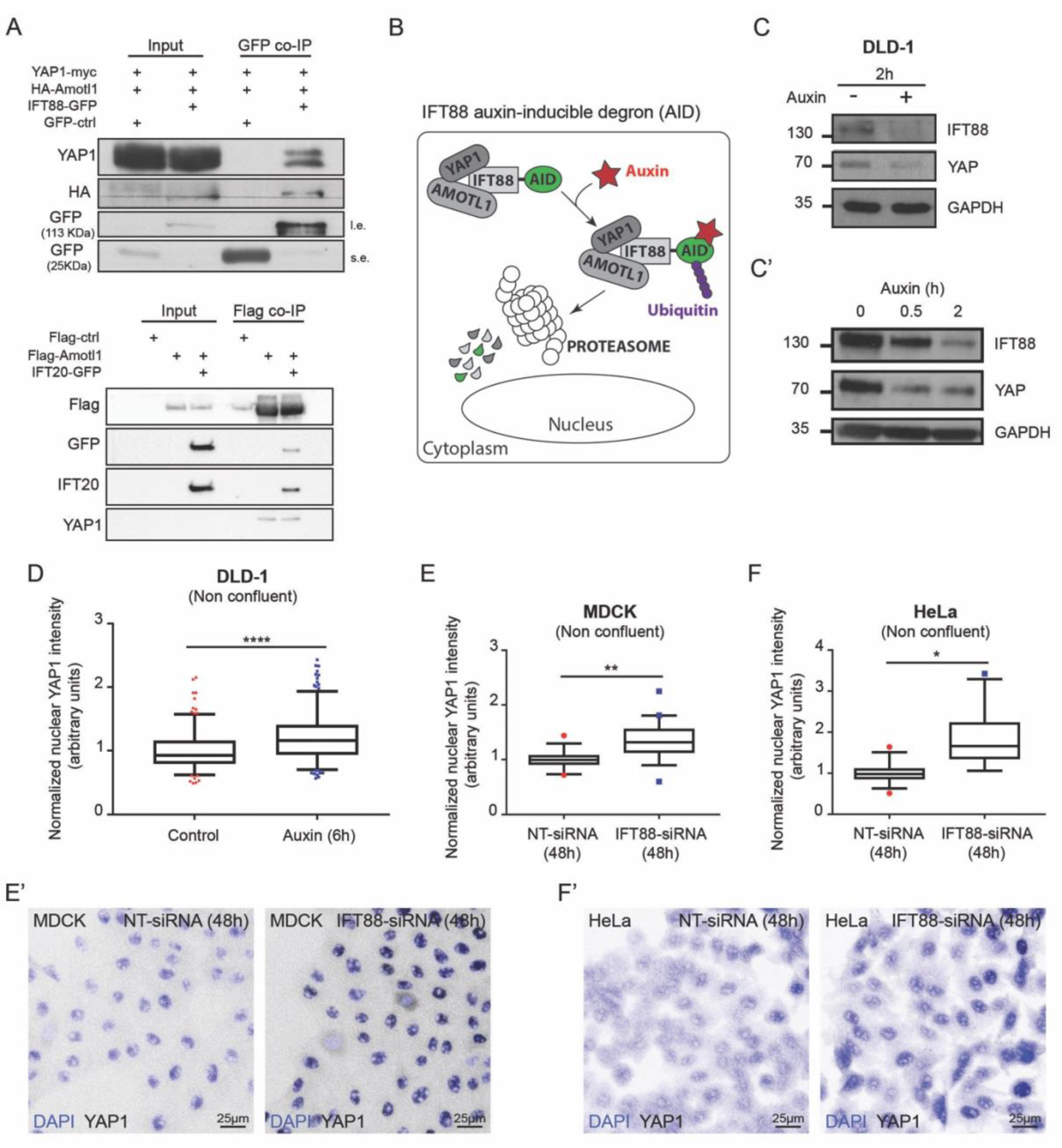
IFT88 and IFT20 are physically associated to YAP1 and IFT88 modulates YAP1 activity. **(A)** Co-IP experiment using HeLa cells transfected with IFT88-GFP, HA-Amotl1 and YAP1-Myc. (l.e. = long exposure. s.e. = short exposure). Co-IP experiment using HEK293 cells transfected with Flag-Amotl1 and IFT20-GFP. Endogenous levels of Yap1 are monitored. **(B)** Schematic representation of the IFT88 auxin-inducible degron (AID) system **(C)** Western blot analysis of IFT88 AID DLD-1 cells after 2 hours auxin treatment. **(C’)** Western blot analysis of IFT88 and YAP1 degradation after auxin treatment (0 h, 0.5h and 2 h). **(D)** Graph shows the increase in normalized YAP1 nuclear signal in cells treated with auxin (6h) (Mann-Whitney p-value <0.0001). (controls: 2 replicates, n=204 cells; Auxin 2h: 2 replicates, n=261 cells). **(E)** Graph shows the increase in YAP/WWTR1 (TAZ) nuclear signal in IFT88-siRNA (48h) treated MDCK cells (n=5 replicates, average cell number analyzed for each condition = 382, t-test p-value 0.004). Box and whiskers (5-95 percentile). Outliers are represented as red dots (NT-siRNA) or blue squares (IFT88-siRNA). **(E’)** Immunofluorescence confocal images (z-projection) of MDCK cells treated with NT – or IFT88-siRNA (48h). DAPI (blue) and YAP/WWTR1 (TAZ) (white inverted LUT). **(F)** Graph shows the increase in YAP/WWTR1 (TAZ) nuclear signal in IFT88-siRNA (48h) treated HeLa cells (n=5 replicates, average cell number analyzed for each condition = 252, t-test p-value 0.02). Box and whiskers (5-95 percentile). Outliers are represented as red dots (NT-siRNA) or blue squares (IFT88-siRNA). **(F’)** Immunofluorescence confocal images (z-projection) of HeLa cells treated with NT – or IFT88-siRNA (48h). DAPI (blue) and YAP/WWTR1 (TAZ) (white inverted LUT).

To functionally assess the impact of endogenous IFT88 depletion on YAP1 activity, we next used a DLD-1 IFT88-auxin-inducible degron (AID) cell line, in which a rapid degradation of IFT88 protein can be induced by auxin treatment (**Fig. 4 B**). Of note, DLD-1 cells don’t grow cilia allowing us to explore the cytoplasmic, cilia independent function of IFT88 (64).

We observed that 2 hours of auxin treatment led to IFT88 and YAP1 degradation. The rapid co-degradation further suggested that both proteins physically interact (**Fig. 4 C**). We confirmed this finding by performing shorter auxin treatments to study the progressive degradation of IFT88 and YAP1 (**Fig. 4 C’**). Importantly, as previously observed *in vivo*, longer depletion of endogenous IFT88 after 6 hours of auxin treatment led to an increase in nuclear YAP1 measured by immunofluorescence (**Fig. 4 D**) (**Supp. 8 A**). Consistently, the nuclear/cytoplasmic YAP1 ratio was also increased (**Supp. 8 A**). Additionally, we confirmed these results using HeLa and MDCK cell lines where IFT88 function was inactivated by IFT88-siRNA. The IFT88-siRNA efficiency was validated by immunofluorescence and western blot (**Supp. 9 A-C**). After 48 hours of IFT88-siRNA treatment, nuclear YAP1 signal was measured by immunofluorescence (**Fig. 4 E-F’**). The inactivation of IFT88 was accompanied with increased nuclear YAP1 (**Fig. 4 E, F**). Altogether, these data indicate that IFT88 can interact with YAP1 in the cytoplasm and is involved in modulating the activation of the Hippo pathway effector YAP1.

To confirm that increased nuclear YAP1 is specific of IFT protein function, we performed IFT88 and IFT20 overexpression assays. While the transfection of pEGFP C1 did not alter the nuclear signal between GFP-positive and GFP-negative cells (endogenous control) (**Supp. 6 D**) (**Supp. 7 B-D**), IFT88-GFP and IFT20-GFP overexpression caused a decrease in nuclear signal (**Supp. 7 C, D**). Together, these data confirm that IFT proteins modulate YAP1 activation.

## Discussion

Primary cilia function has long been implicated in the control of the developmental program, but whether cilia-related proteins could affect cell signaling independently of their ciliary function remains unclear. Here, we have shown that IFT complex B members, IFT88 and IFT20 modulate the Hippo pathway effector YAP1 in different cell types. We also demonstrated that IFT88 and IFT20 modulates PE size in mouse and that Ift88 and Ift54 do the same in zebrafish PE and myocardium. Mechanistically, work *in cellulo* suggest that IFT88 physically interacts with YAP1 to modulate its activity. Together, this work establishes a previously unknown role for IFT complex B proteins in regulating cell signaling and tissue growth during cardiogenesis and shed a novel light on cilia related protein function during cardiogenesis.

IFT proteins have long been associated with ciliary functions in developmental processes. For example, *ift88* mutants display a number of phenotypes reminiscent of ciliary defects such as abnormal patterning of the neural tube, defects in the Hedgehog pathway and left-right patterning (58). Ift88 has also been associated with planar cell polarity (65) and cell division (15, 16). Our observations that the *ift88* mutants show increased number of nuclear Yap1-positive cells in the myocardium and that the *Ift20 KO* mice display increased nuclear Yap1/Amot1 – positive cells in the PE, provide evidences for a novel role of IFT complex B proteins in cardiogenesis. Mechanistically, IFT proteins are best known for their function in ciliary transport (66). Here, we describe an unexpected interaction between the ciliary machinery proteins and a potent mechanosensing pathway, the Hippo pathway. The hippo effector YAP1 is known to have essential roles in cancer (67), regeneration (24,25,68), and organ size control (69–71). Several mechanisms have been shown to regulate the shuttling of YAP1 into the nucleus, including phosphorylation by Hippo kinases. Recent studies show that YAP1 is mechanosensitive and that force applied to the nucleus can directly drive YAP1 nuclear translocation (27, 72). Additionally, Angiomotin (AMOT) have been shown to interact physically with YAP1 and acts as a buffering factor sequestering YAP1 in the cytoplasm (31). Nevertheless, AMOTL1 has also been shown to co-localize with YAP1 in the nucleus (23, 73) demonstrating YAP1 subcellular localization is highly regulated by Motin family proteins. Our results indicate that IFT complex B proteins are also involved in regulating YAP1 localization. We demonstrate that IFT88 interacts biochemically with YAP1 and both co-localize in the cytoplasm. We did not study TAZ, the other Hippo effector which is known to act with YAP1 (74) and we cannot draw general conclusions on the role of IFT88 on all the known Hippo effectors. Nevertheless, our work suggests alternative ways to interpret Ift88 mutant phenotypes, which are often interpreted based on polarity or cilia function issues, and, more generally, phenotypes of other mutants with abnormal IFT complex B proteins. Our working model is that Ift88 participates in sequestering YAP1 away from the nucleus using its cargo transport activity. Other cilia related proteins such kinesin2 and IFT complex A proteins have been shown to promote nuclear localization of β-catenin during Wnt signaling in drosophila (75), further suggesting that proteins identified for their ciliary transport functions are not always limited to that function. Interestingly, IFT-A complexes control retrograde protein transport, from the tip to the base of the cilium (76–78), while the IFT-B complexes do the opposite, as observed for YAP1 transport towards the nucleus. Future work will reveal the mechanism by which Ift88 limits nuclear translocation of YAP1 in this process and address whether IFT complex A proteins play a role.

Importantly, while our study points towards a non-canonical function for IFT complex B proteins, our results do not exclude a role for primary cilia in PE formation. We found that *iguana/dzip* mutants display a avcPE phenotype suggesting that primary cilia function is required for PE morphogenesis. The HH pathway is often associated with ciliary function. To date, a number of studies suggest the proepicardium and epicardium formation are not regulated by Hedgehog signaling (79, 80). Indeed, dissection of zebrafish *shha* function in the PE and epicardium using a *tcf21:CreER*, a well-established PE and epicardial tissue driver (81), does not affect PE formation (80). Besides, expression of *Shh*, *Dhh*, *Ihh*, and *Ptch1* was neither detected in the mouse PE nor in the epicardium at subsequent stages (79). Thus, primary cilia certainly operate independently of the HH pathway in the process. Our work further highlights the important role of the BMP pathway in PE formation. BMP is essential for PE formation and morphogenesis (50, 82). There is numerous evidence to suggest that primary cilia modulate BMP pathway and, more generally, TGF beta signaling pathways (83, 84). In endothelial cells, primary cilia modulate angiogenesis by altering BMP signaling in endothelial cells (85). We speculate the situation is different in PE cells, where BMP is activated downstream of YAP1, and where IFT88 helps to limit YAP1 and BMP signaling. Similarly, secondary heart field development also depends on the modulation of Hippo signaling by BMPs (21).

In summary, our study is the first report implicating IFT complex B proteins during PE development by modulating YAP1 activity independently of any cilia function. A deeper understanding of the molecular mechanisms regulating PE development is of great importance, underlined by its role in myocardial maturation and coronary vessel formation during development, as well as the regenerative capacity of the heart (86). Considering that IFT proteins are often associated with ciliopathies, linking IFT with YAP1 activity might have important implications for understanding the etiology of ciliopathies during cardiogenesis and for the interpretation of ciliary defects in IFT mutants.

## Acknowledgements

We thank P. Lamperti, E. Steed, A Bhat, D. Riveline, S. Harlepp, J. Godin, A. Benmerah and the Vermot lab for discussion and thoughtful comments on the manuscript, in particular R. Chow for her help with editing. We thank G. Pazour, M. Faucourt and N. Spassky for advice, the gift of plasmids and for providing the *Ift20* and *Ift88* mutant mouse lines and C. Cimper, L. Bombardelli and C. Shea for technical assistance. We are grateful to the IGBMC fish facility, the IGBMC imaging center and the imaging platforms of the SFR Necker. This project has received funding from the European Union’s Horizon 2020 research and innovation programme under the Marie Sklodowska-Curie grant agreement No 708312 (MP) and from the European Research Council (ERC) under the European Union’s Horizon 2020 research and innovation programme: GA N°682938 (JV). This work was supported by FRM (DEQ20140329553), ANR (ANR-15-CE13-0015 – liveheart) and by the grant ANR-10-LABX-0030-INRT, a French State fund managed by the Agence Nationale de la Recherche under the frame program Investissements d’Avenir labeled ANR-10-IDEX-0002–02. BD team was supported by ANR-12-CHEX-005 and CNRS. SM team was supported by core funding from the Institut *Imagine*, the Institut Pasteur, the INSERM, the Université Paris Descartes, and a grant from the AFM-Téléthon [Trampoline 18727]. TL was funded by the ED515 (1691/2014). L.L. is supported by the European Commission (H2020-MSCA-ITN-2016 European Industrial Doctorate 4DHeart 722427).

---

**Supplementary Figure 1.**
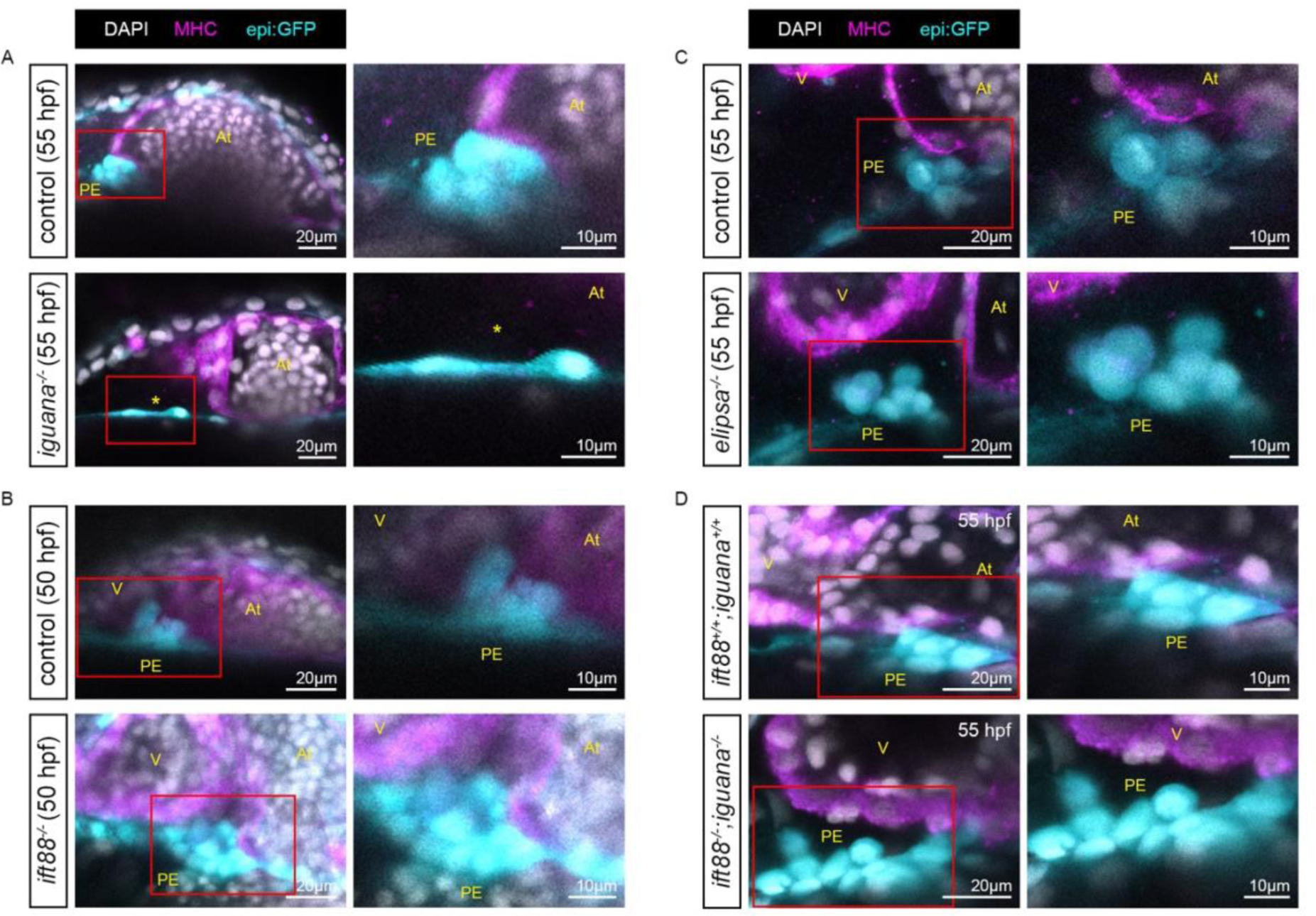
IFT complex B mutants show increased proepicardial size in zebrafish. **(A)** Confocal sections of whole mount immunofluorescence of iguana-/-, epi:GFP and control using anti-myosin heavy chain antibody (MHC) (magenta), anti-GFP (cyan) and DAPI (white) antibodies at 55 hpf. Asterisk shows lack of avcPE as there is only one rounded PE cell. **(B)** Confocal sections of whole mount immunofluorescence of ift88-/-, epi:GFP and control using anti-myosin heavy chain antibody (MHC) (magenta), anti-GFP antibody (cyan) and DAPI (white) at 50 hpf. **(C)** Confocal sections of whole mount immunofluorescence of elipsa-/-, epi:GFP and control using anti-myosin heavy chain antibody (MHC) (magenta), anti-GFP (cyan) and DAPI (white) antibodies at 55 hpf. **(D)** Confocal sections of whole mount immunofluorescence of ift88-/-, iguana-/-, epi:GFP and ift88+/+, iguana+/+, epi:GFP using anti-myosin heavy chain antibody (MHC) (magenta), anti-GFP (cyan) and DAPI (white) antibodies at 55 hpf. All images are ventral views, anterior is to the top. V, ventricle; At, atrium; PE, avcPE. Zoomed regions contained inside the red boxes are showed on the right side.

**Supplementary Figure 2.**
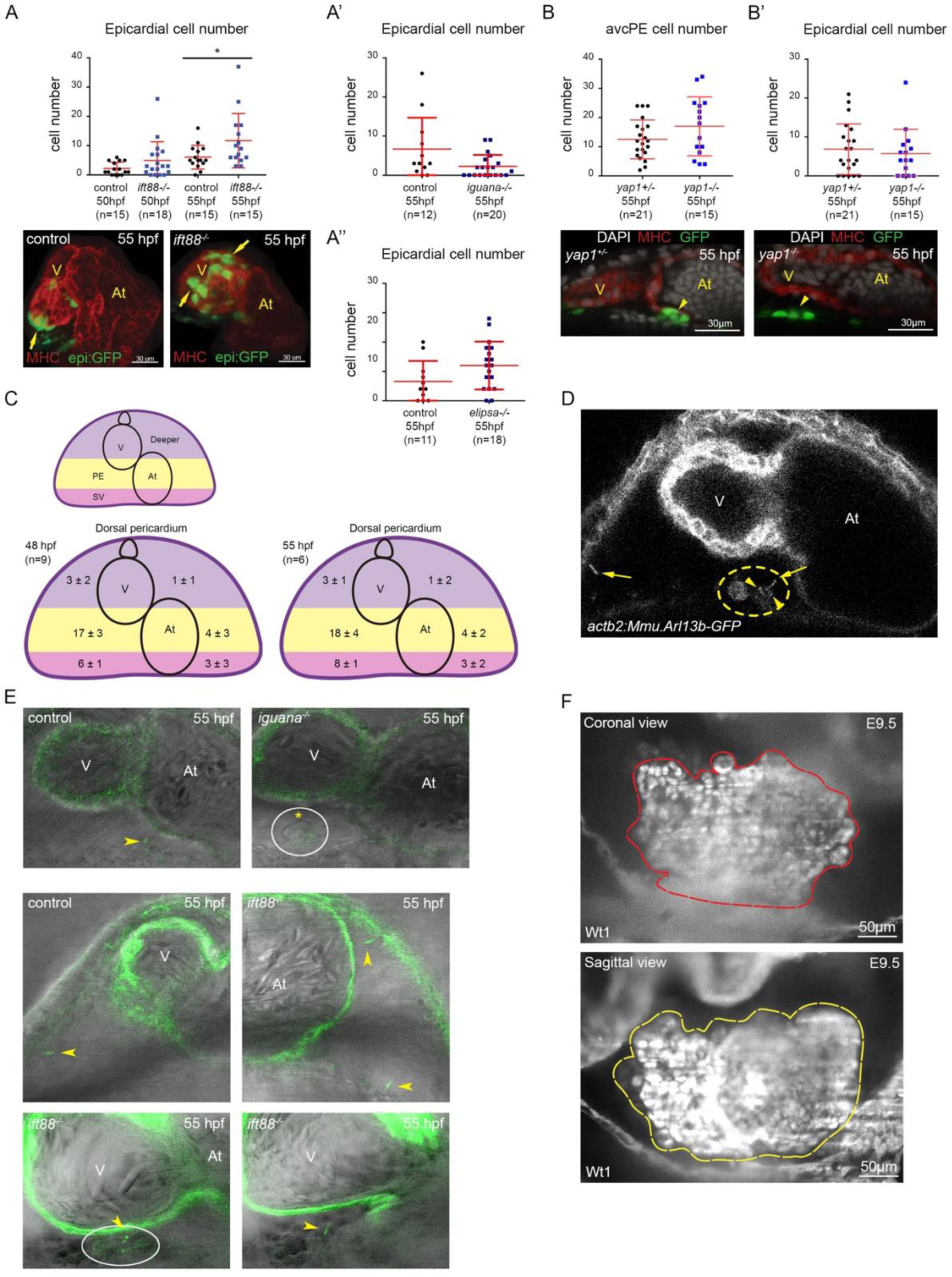
Cilia protruding into the pericardial cavity are distributed heterogeneously in zebrafish during proepicardial development. **(A-A’’)** Graphs show epicardial cell numbers quantified in ift88, iguana and elipsa mutants in epi:GFP background. **(A)** At 55 hpf, ift88 mutants (n=15) showed increased epicardial cell numbers (t-test p value 0.04). 3D projections of whole mount immunofluorescence of hearts using anti-myosin heavy chain antibody (red) and GFP (green) expression. Ventral view, anterior is to the top. Arrows mark some epicardial cells; (**A’**) iguana mutants (n=20) showed a tendency towards decreased epicardial cell numbers (t-test p value 0.055), while (**A’’**) elipsa mutants (n=18) showed a tendency towards increased epicardial cell number (t-test p value 0.08). **(B)** Graph shows avcPE cell numbers quantified in yap1-/- (n=15) and control (n=21) embryos in tcf21:nsl-GFP background at 55 hpf. (t-test p-value 0.12) Control and yap1-/- immunofluorescence confocal sections labelled with anti-myosin heavy chain antibody (MHC) (red), GFP (green), DAPI (white). Ventral view, anterior is to the top. Yellow arrowheads point at the avcPE. **(B’)** Graph shows epicardial cell numbers quantified in yap1-/- (n=15) and control (n=21) embryos in tcf21:nsl-GFP background at 55 hpf. (Mann Whitney p-value 0.63) **(C)** For cilia quantification, we divided the dorsal pericardial wall in to three different regions: SV region, including the sinus venosus (pink); PE region, where the avcPE forms (yellow) and Deeper region (purple). These three regions were subdivided in to right and left halves, containing the ventricle or the atrium respectively. At 48 hpf (n=9 larvae), prior to PE formation, cilia protruding from the dorsal pericardium showed a heterogeneous distribution. Interestingly, the right half of the SV (6 ± 1) and the PE (17 ± 3) regions, where both PE clusters will form, presented higher cilia number than the rest of the regions. At 55 hpf, when the avcPE is formed, the cilia distribution was similar to that observed at 48 hpf (n= 6 larvae). **(D)** Confocal section of actb2:Mmu.Arl13b-GFP embryo (55 hpf). Yellow dotted circle encloses the avcPE. Yellow arrows point to cilia protruding from the ventral and dorsal pericardium. Yellow arrowheads point at immotile and bent cilia protruding from a few avcPE cells. **(E)** Confocal section of iguana; actb2:Mmu.Arl13b-GFP and control embryos (55 hpf). Yellow arrowhead points at cilium protruding from the avcPE. Yellow asterisk shows lack of cilia in the avcPE (enclosed in the white circle). Confocal section of ift88; actb2:Mmu.Arl13b-GFP and control embryos (55 hpf). In control embryo, yellow arrowhead points at cilium protruding from the ventral pericardium. In ift88 mutant embryo, yellow arrowheads point at cilia protruding from ventral and dorsal pericardium and the avcPE (enclosed in the white circle). **(F)** Coronal and sagittal sections acquired by light sheet microscopy to illustrate the methods used to measure PE volume (labeled with anti-Wt1 antibody). Red and yellow dotted shapes enclose the PE area. In all images ventral views, anterior is to the top. V, ventricle; At, atrium. In all graphs, red bars indicate mean ± standard deviation.

**Supplementary Figure 3.**
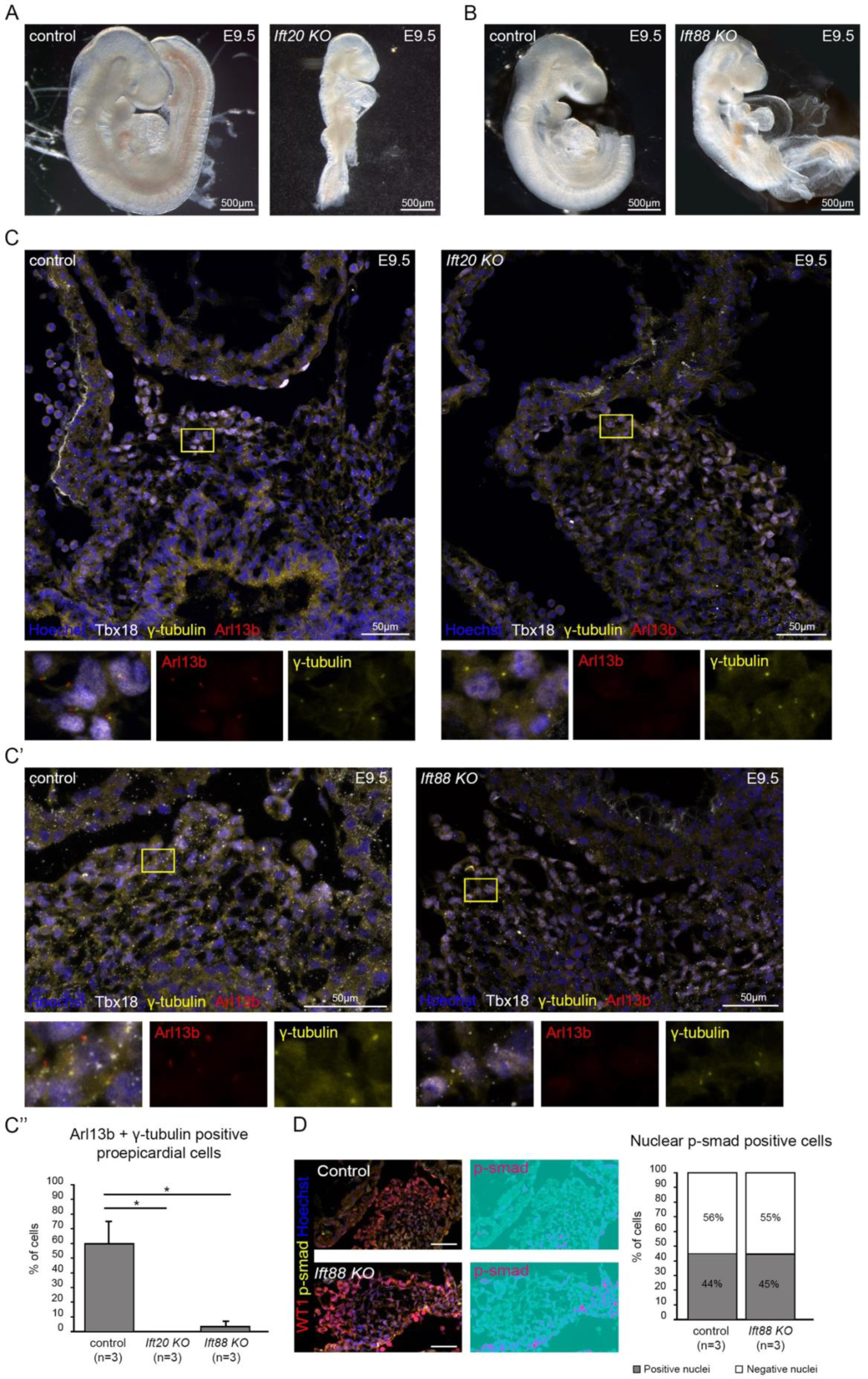
Ift20 KO mice proepicardial cells lack cilia. **(A,B)** Ift20 and Ift88 KO mice show left-right patterning defects including heart looping defects at E9.5. **(C)** Control and Ift20 KO cryosections imaged by confocal microscopy after labelling with anti-TBX18 (white), anti-Arl13b (red) and anti-γ-tubulin (yellow) antibodies and Hoechst (blue) at E9.5. Zoomed region (enclosed in yellow box) shows the lack of Arl13b signal in Ift20 KO mice. Individual channels are shown for Arl13b (red), γ-tubulin (yellow). **(C’)** Control and Ift88 KO cryosections labelled with anti-TBX18 (white), anti-Arl13b (red) and anti-γ-tubulin (yellow) antibodies and Hoechst (blue) at E9.5. Zoomed region (enclosed in yellow box) shows the decrease of Arl13b signal in Ift88 KO mice. Individual channels are shown for Arl13b (red), γ-tubulin (yellow). **(C’’)** Graph shows the percentage of ciliated PE cells in Ift20 KO (n=3, Ift88 KO (n=3) and control (n=3) mice. The percentage of ciliated PE cells is severely reduced in Ift20 KO (n=3) and Ift88 KO (n=3) when compared to control (n=3) mice. (t-test Ift20 p-value 0.024; t-test Ift88 p-value 0.025) **(D)** Control and Ift88 KO cryosections labelled with WT1 (red), p-smad 1/5/9 (yellow) and Hoechst (blue) at E9.5. Individual channel is displayed for p-smad 1/5/9 as ice LUT to facilitate the visualization of signal intensity (green is the minimum and red is the maximum).Graph shows that the percentage of p-smad 1/5/9 positive PE cells is similar in Ift88 KO (n=3 embryos: 1477 nuclei analyzed) and controls (n=3 embryos: 1514 nuclei analyzed) (Chi-square test of homogeneity =0.15225, p-value 0.6964 on 1 degree of freedom).

**Supplementary Figure 4.**
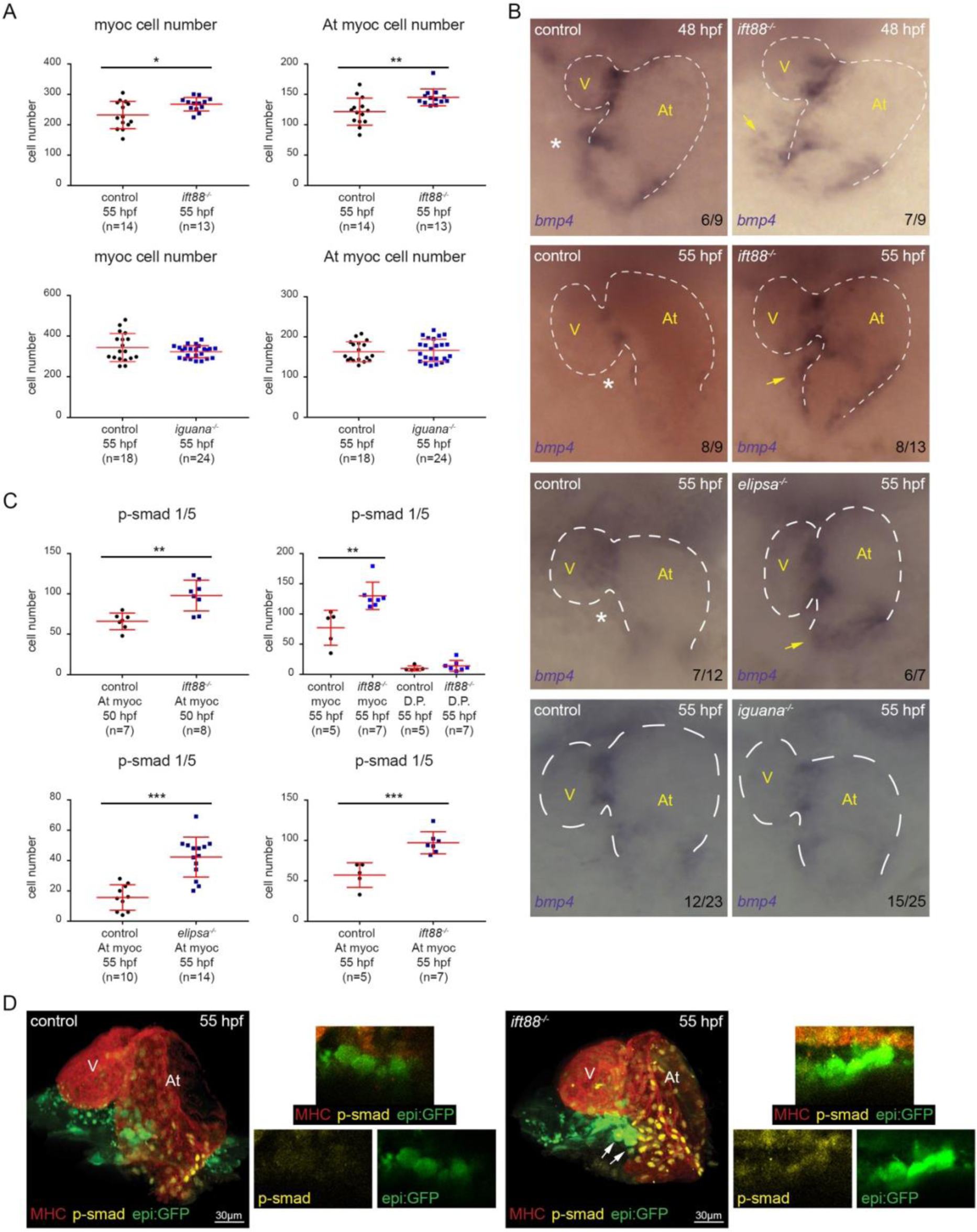
bmp4 is overexpressed in ift88, elipsa/ift54 and Ift20 mutants. **(A)** The top two graphs show total myocardial and atrial-myocardial cell numbers quantified in ift88 (n=13) mutants and controls (n=14) at 55 hpf. (t-test total myocardium p-value 0.016; atrial myocardium p-value 0.003). The bottom two graphs show total myocardial and atrial-myocardial cell number quantified in iguana (n=24) mutants and controls (n=18) at 55 hpf. (t-test total myocardium p-value 0.2; atrial myocardium p-value 0.68). **(B)** Whole mount bmp4 in situ hybridization performed on control and ift88 mutant embryos at 48 hpf and at 55 hpf on ift88, elipsa and iguana mutants and their controls. Yellow arrows point to bmp4 overexpression, while white asterisks mark reduced or absent expression. Ventral views, anterior is to the top. V, ventricle; At, atrium. **(C)** Graphs show number of p-smad1/5 positive cells in the atrial myocardium quantified in ift88 (at 50 hpf n=8; at 55 hpf n=7), elipsa (n=14) mutants and their controls (n=7; n=5; n=10 respectively). At 50 hpf, ift88 mutants show p-smad 1/5 increased cell number on the atrial myocardium (t-test p value 0.0017). Similar data were obtained at 55 hpf (t-test p value 0.0008). At 55 hpf, elipsa mutants also show p-smad 1/5 increased cell number in the atrial myocardium (t-test p value 0.0003). At 55 hpf, ift88 mutants show p-smad 1/5 increased cell numbers in the myocardium (t-test p value 0.005). **(D)** 3D projections of whole mount immunofluorescence of hearts using anti-myosin heavy chain antibody (MHC) (red), epi:GFP (green) and anti-p-smad1/5 (yellow) antibody. Arrows mark avcPE cells positive for epi:GFP and p-smad1/5. Zoomed confocal sections show avcPE in ift88 mutant and control embryos. Individual channels are displayed for p-smad 1/5 and GFP. Ventral views, anterior is to the top. In all graphs, red bars indicate mean ±standard deviation.

**Supplementary Figure 5.**
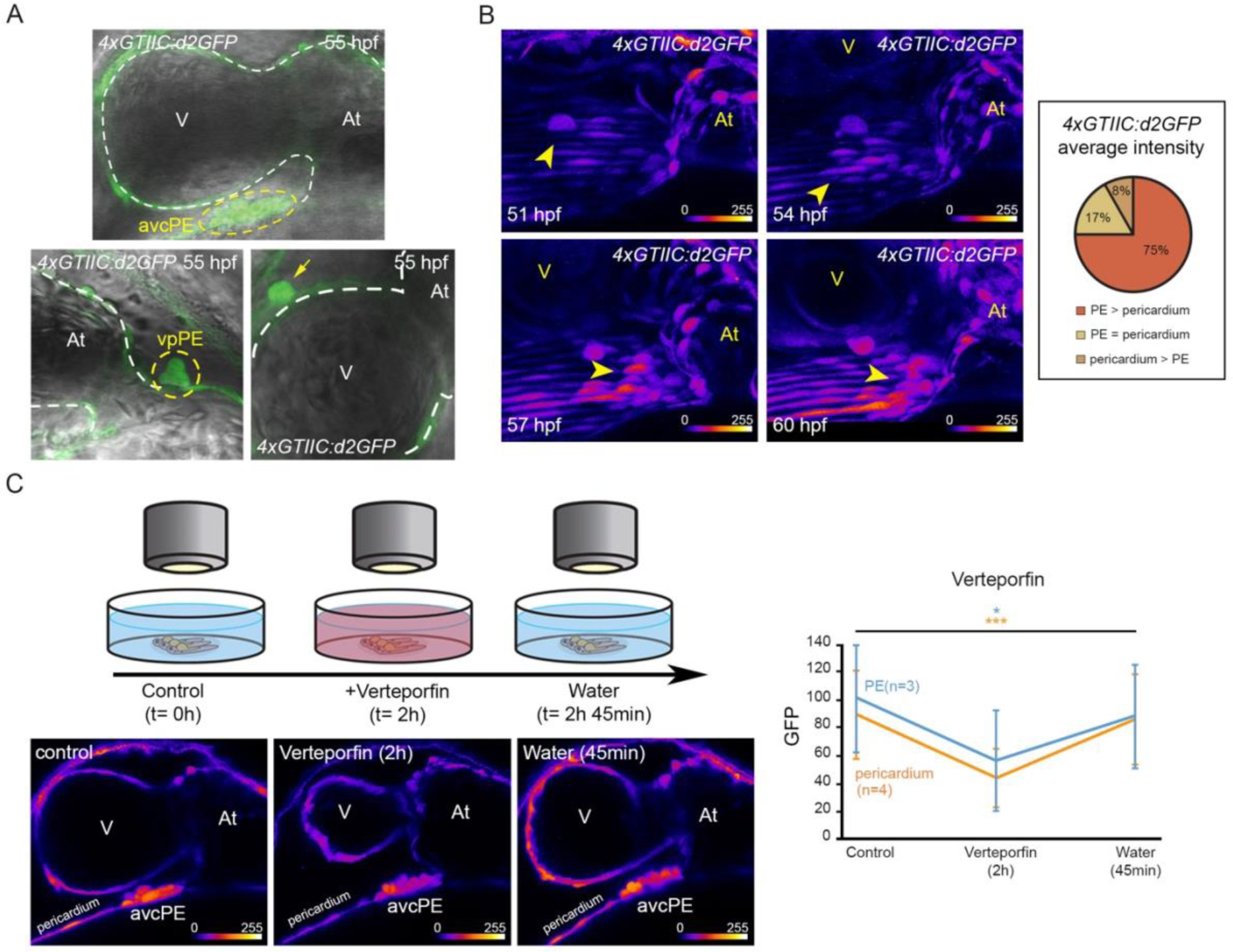
Yap1-Tead activity during proepicardial development in zebrafish. **(A)** Confocal sections of 4xGTIIC:d2GFP signal merged with bright-field, showing Yap/Wwtr1-Tead interaction in avcPE, vpPE, epicardial, myocardial and pericardial cells (n=12). Yellow dashed circles enclose the avcPE and the vpPE. Arrow shows an epicardial cell. Ventral views, anterior is to the top. **(B)** Maximum projection 4xGTIIC:d2GFP time lapse (51-60 hpf) snapshots (n=6). Arrowheads point at the avcPE. GFP signal is shown as fire LUT where blue is the minimum and yellow is the maximum to facilitate visualization of the intensity changes through the experiment. Ventral views, anterior is to the top. V, ventricle; At, atrium. Graph shows 75% of the embryos displayed higher average GFP intensity in PE cells than in pericardial cells (n=12, 7-15 cells of each type). **(C)** Scheme of the experiment to assess Verteporfin specificity. Time lapse performed on 4xGTIIC:d2GFP embryos at 55 hpf. Yap/Wwtr1-Tead activity (average GFP intensity) was measured on the same PE (3 cells in each embryo, n=3) and pericardial (3 cells in each embryo, n=4) cells at three timepoints: before adding Verteporfin (5µM) (t=0h), after 2 hours of treatment (t=2h) and 45 min after washing out the Verteporfin with fish water (t=2h45 min). Graph shows the decrease in Yap/Wwtr1-Tead activity (average GFP intensity) due to Verteporfin treatment on PE (blue) and pericardial (orange) cells, which is rescued after removing the inhibitor. (Pericardial cells ANOVA p value 0.0007. PE cells ANOVA p value 0.05). Example of 4xGTIIC:d2GFP embryo confocal sections used for the experiment. GFP signal is shown as fire LUT where blue is the minimum and yellow is the maximum to facilitate visualization of the intensity changes through the experiment. Ventral views, anterior is to the top. V, ventricle; At, atrium.

**Supplementary Figure 6.**
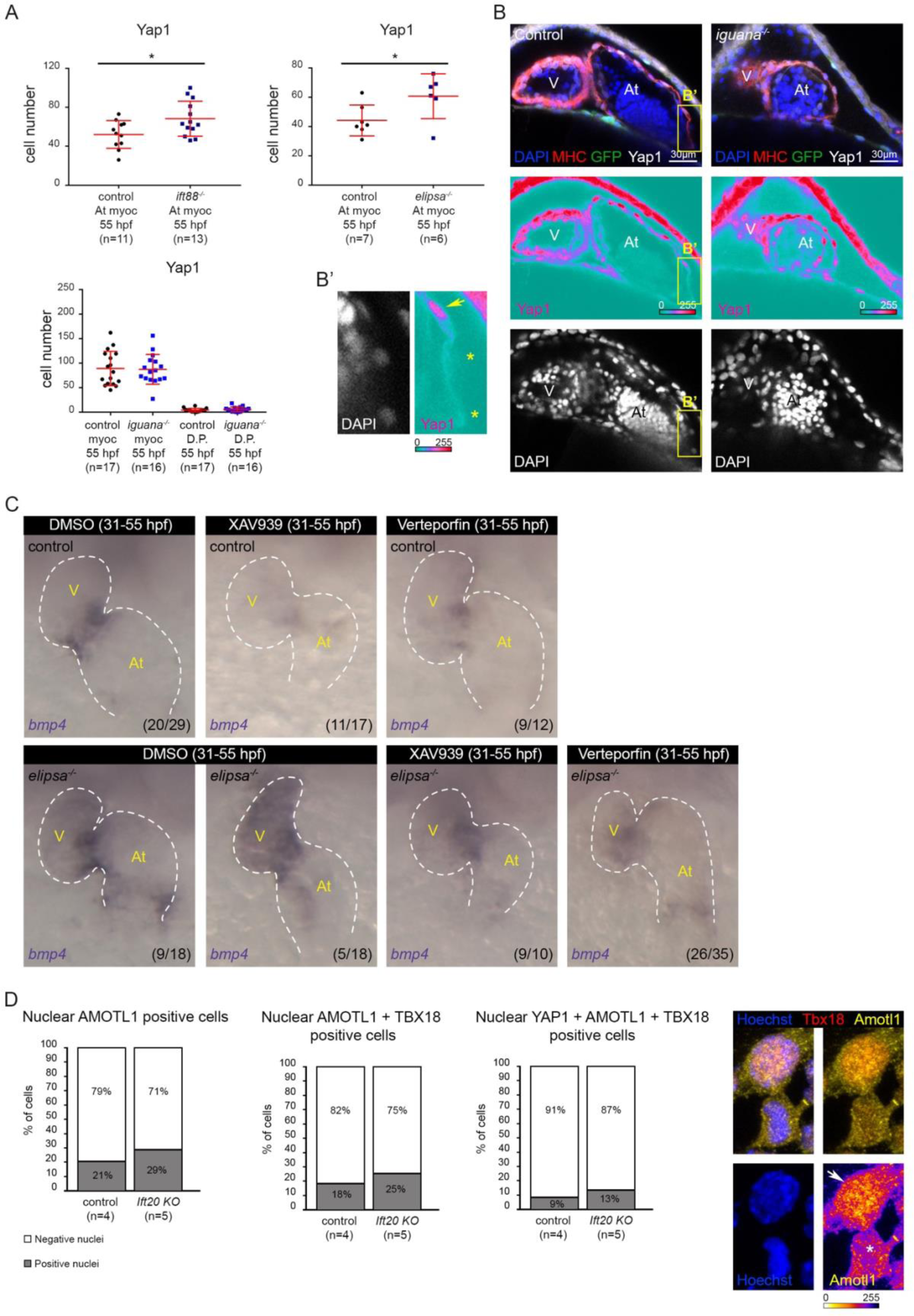
Yap1 activity is increased in the myocardium of ift88 and elipsa/ift54 zebrafish mutants and AMOTL1 activity is increased in IFT20 KO mice proepicardium. **(A)** The top two graphs show number of Yap1-positive cells in the atrial myocardium quantified in ift88-/-, epi:GFP (n=13) and elipsa-/-, epi:GFP (n=6) mutants and their controls (n=11 and n=7 respectively) at 55 hpf. Mutants show increased Yap1-positive cell numbers (t-test ift88 p value 0.024 and elipsa p value 0.042). Bottom graph shows number of Yap1-positive cells in the myocardium and dorsal pericardium (D.P.) quantified in iguana-/-, epi:GFP (n=16) mutants and their controls (n=17) at 55 hpf (t-test myocardium p value 0.875 and D.P. p value 0.312). **(B)** Control and iguana-/-, epi:GFP immunofluorescence confocal sections labelled with anti-myosin heavy chain antibody (MHC) (red), GFP (green), anti-Yap1 antibody (white) and DAPI (blue) at 55 hpf. Individual channel is displayed for Yap1 (signal is shown as ice LUT to facilitate visualization of signal intensity, where green is the minimum and red is the maximum) and DAPI (white). Ventral view, anterior is to the top. **(B’)** Zoomed region (yellow box in panel **B**) shows Yap1 and DAPI channels to illustrate the method used to quantify Yap1-positive (Yap1 signal in the nucleus: yellow arrow) and –negative (yellow asterisks) cells. **(C)** Whole mount bmp4 in situ hybridization in untreated elipsa mutant (n=18) and control (n=29) embryos and treated with XAV939 (10µM) (elipsa mutant, n=10 and control, n=17) or Verteporfin (20µM) (elipsa mutant, n=35 and control, n=12) from 31 to 55 hpf. Treated embryos showed either decreased or absent bmp4 expression at the atrioventricular canal myocardium and the venous pole. Ventral views, anterior is to the top. **(D)** Graphs show the percentages of AMOTL1-positive PE cells, double AMOTL1-TBX18-positive PE cells and triple YAP1-AMOTL1-TBX18-positive PE cells in Ift20 KO (n=5 embryos: 1196 nuclei analyzed) and control (n=4 embryos: 929 nuclei analyzed) mice at E9.5. The percentage of nuclear AMOTL1-positive cells (Chi-square test of homogeneity = 14,748, p-value 1,23E-04 on 1 degree of freedom), nuclear AMOTL1-TBX18-positive cells (Chi-square test of homogeneity = 12,506, p-value 4,06E-04 on 1 degree of freedom) and nuclear YAP1-AMOTL1-TBX18-positive cells (Chi-square test of homogeneity = 6,9059, p-value 8,59E-03 on 1 degree of freedom) were higher in Ift20 KO than in control mice. Control (n=4) and Ift20 KO (n=5) sections labelled with TBX18 (red), Amotl1 (yellow) and Hoechst (blue). Individual AMOTL1 channel shows the difference between nuclear AMOTL1-positive cells (white arrow) and AMOTL1-negative cells (white asterisk). AMOTL1 signal is shown as fire LUT to facilitate visualization of the signal intensity, where blue is the minimum and yellow is the maximum. In all graphs, red bars indicate mean ±standard deviation. V, ventricle; At, atrium.

**Supplementary Figure 7.**
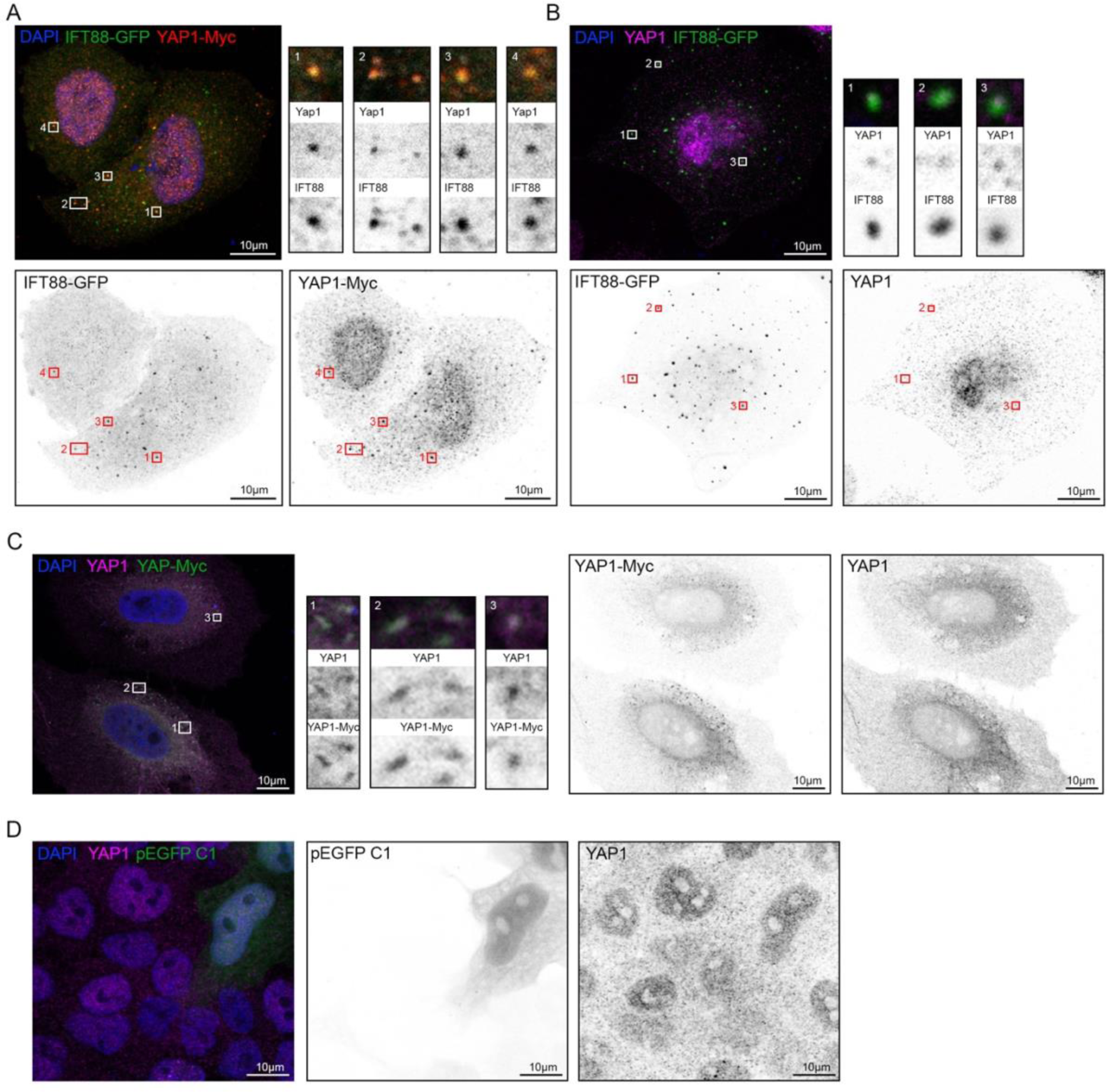
IFT88-GFP co-localize with YAP1 in the cytoplasm. **(A)** Confocal section of HeLa cells transfected with IFT88-GFP and Yap1-Myc plasmids (48h). DAPI (blue), IFT88-GFP (green) and Yap1 (visualized using anti-Myc antibody) (red). Individual channels are displayed for IFT88-GFP and Yap1-Myc. (**1-4**) Zoom of selected areas inside boxes showing co-localization. **(B)** Confocal sections of HeLa cells transfected with IFT88-GFP (48h) showing co-localization with endogenous YAP (visualized using anti-YAP/WWTR1 (TAZ) antibody). DAPI (blue), IFT88-GFP (green) and YAP1 (magenta). (**1-3**) Zoom of the selected areas inside white boxes showing co-localization of IFT88 and YAP1. Individual channels are displayed for IFT88-GFP and YAP1 signals. **(C)** Confocal sections of HeLa cells transfected with Yap1-Myc (48h) showing co-localization with endogenous YAP1 (visualized using anti-YAP/WWTR1 (TAZ) antibody). DAPI (blue), Yap1-Myc (green) and YAP1 (magenta). (**1-3**) Zoom of the selected areas inside white boxes showing examples of co-localization between Yap1-Myc (visualized using anti-Myc antibody) and endogenous YAP1 (using YAP/WWTR1 (TAZ) antibody) signals. Individual channels are displayed for Yap1-Myc and YAP1 signals. **(D)** Confocal sections of HeLa cells transfected with pEGFP C1 (48h) do not show co-localization with endogenous YAP1 (visualized using anti-YAP/WWTR1 (TAZ) antibody). DAPI (blue), pEGFP C1 (green) and YAP1 (magenta). Individual channels are displayed for pEGFP C1 and YAP1 signals.

**Supplementary Figure 8.**
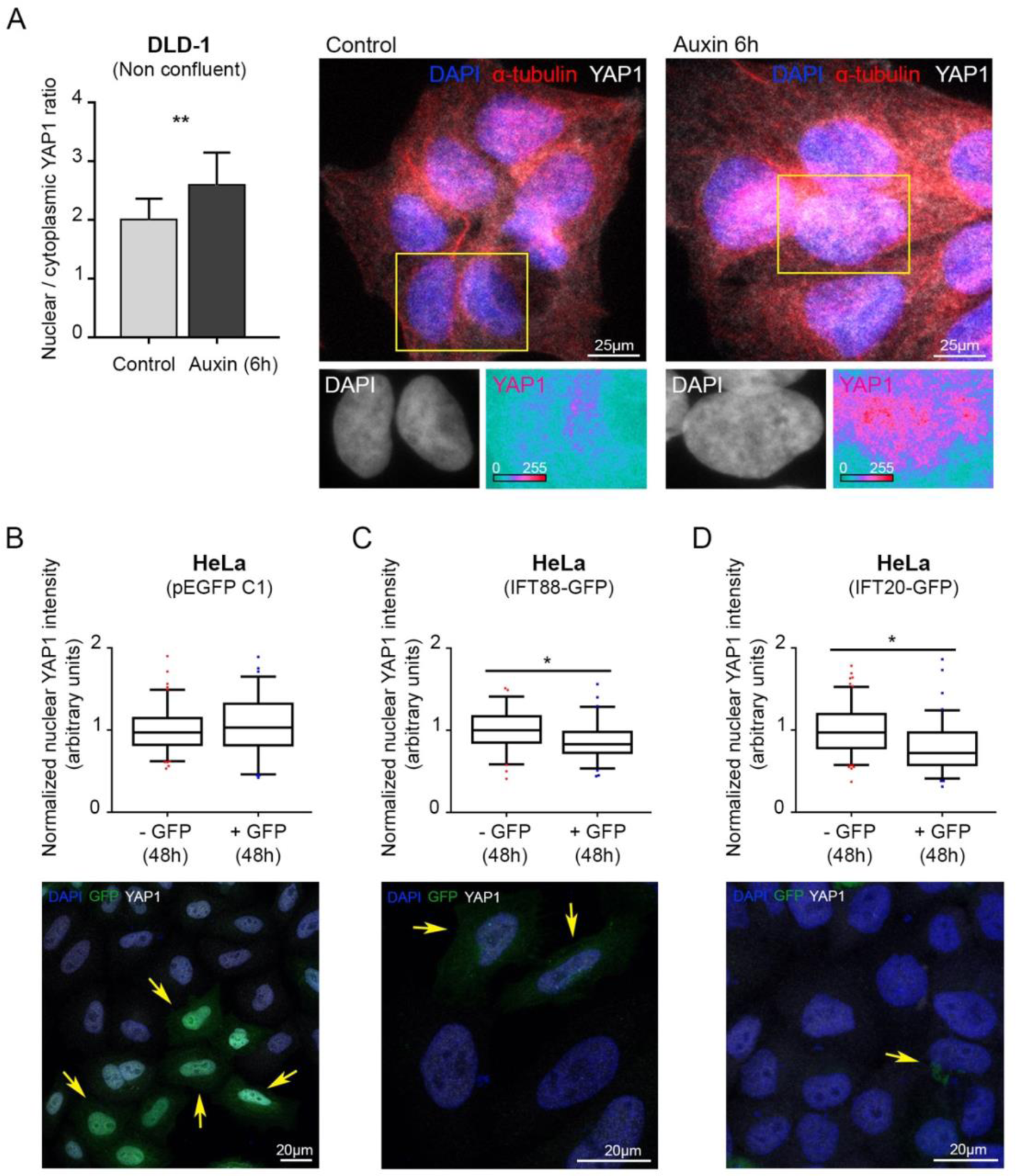
IFT88 and IFT20 regulate YAP1 activity. **(A)** YAP1 Nuclear/cytoplasmic ratio is increased in cells treated with auxin (6h) (Mann-Whitney p-value 0.004). (controls: 2 replicates, n=102 cells; Auxin 6h: 2 replicates, n=104 cells). Immunofluorescence confocal images (z-projection) of DLD-1 cells treated with auxin (6 h) and controls. DAPI (blue), YAP1 (white) and α-tubulin (red). Zoomed regions show DAPI and YAP1 channels. YAP1 channel is shown in ice LUT where green is the minimum and red is the maximum to facilitate the visualization of signal intensity increases after treatment. **(B-D)** IFT88 and IFT20 overexpression assays performed in HeLa cells (48h). Graphs show nuclear YAP/WWTR1 (TAZ) signal in GFP-positive cells (+ GPF), compared to GFP-negative cells (-GPF: endogenous control) in cells transfected with pEGFP C1 (GFP control) (n=4 replicates, average cell number analyzed for each condition = 22, t-test p-value 0.5) **(B)**, IFT88-GFP (n=3 replicates, average cell number analyzed for each condition = 23, t-test p-value 0.01) **(C)** or IFT20-GFP (n=4 replicates, average cell number analyzed for each condition = 27, t-test p-value 0.03) **(D)**. Box and whiskers (5-95 percentile). Outliers are represented as red dots (-GFP) or blue squares (+GFP). Immunofluorescence confocal images (z-projection) of cells transfected with pEGFP C1 (GFP control), IFT88-GFP or IFT20-GFP. DAPI (blue) and YAP/WWTR1 (TAZ) (white). Yellow arrows mark GFP signal.

**Supplementary Figure 9.**
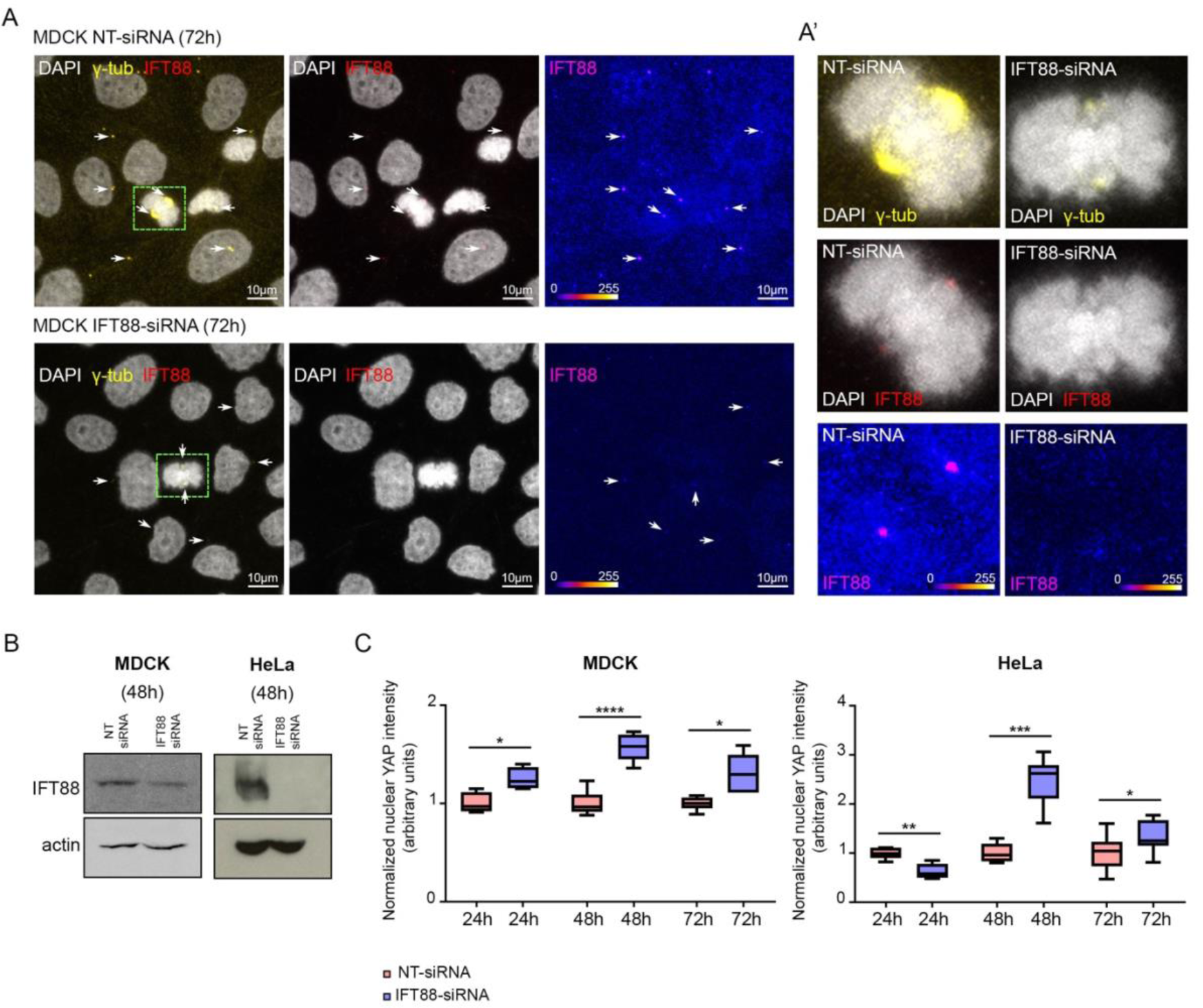
Efficiency validation of IFT88-siRNA in MDCK and HeLa cells. **(A)** Immunofluorescence microscopy images (maximum projection) of MDCK cells treated with NT- or IFT88-siRNA (72h). DAPI (white), γ-tubulin (yellow) and IFT88 (red). White arrows highlight IFT88-positive centrosomes facilitating the visualization of IFT88 signal depletion upon IFT88-siRNA treatment. **(A’)** Zoom of dividing cells (green boxes) treated with NT-siRNA and IFT88-siRNA respectively. Centrosomes show reduced IFT88 (red) and γ-tubulin (yellow) signal after IFT88 depletion. DAPI (white). IFT88 channel is shown in fire LUT where blue is the minimum and yellow is the maximum to facilitate the visualization of the intensity reduction after the treatment. **(B)** Western blot analysis of HeLa and MDCK cells after 48h NT- and IFT88-siRNA treatments respectively. **(C)** Graphs show the increase in nuclear YAP/WWTR1 (TAZ) signal in IFT88-siRNA treated cells (blue), compared to NT-siRNA controls (red) at 24, 48 and 72h. Box and whiskers (5-95 percentile). (MDCK: 24h: n=1 replicate, average cell number analyzed for each condition = 42, t-test p-value 0.02; 48h: n=1 replicate, average cell number analyzed for each condition = 52, t-test p-value <0.0001; 72h: n=1 replicate, average cell number analyzed for each condition = 126, t-test p-value 0.01) (HeLa: 24h: n=1 replicate, average cell number analyzed for each condition = 12, t-test p-value 0.001; 48h: n=1 replicate, average cell number analyzed for each condition = 12, t-test p-value 0.0002; 72h: n=1 replicate, average cell number analyzed for each condition = 32, t-test p-value 0.047).

**Supplementary Table 1.** Yap/Wwtr1-Tead activity (GFP average intensity) in PE and pericardial cells.

| Average intensity | Embryo_01 | Embryo_02 | Embryo_03 | Embryo_04 | Embryo_05 | Embryo_06 |
| --- | --- | --- | --- | --- | --- | --- |
| **PE cells** | 71,37 | 72,42 | 30,84 | 89,57 | 69,72 | 49,63 |
| St-Dev | 11,91 | 21,49 | 26,85 | 20,56 | 24,85 | 13,35 |
| **Pericardial cells** | 52,45 | 45,17 | 55,81 | 94,08 | 49,98 | 29,95 |
| St-Dev | 21,16 | 9,76 | 18,25 | 29,84 | 14,67 | 7,51 |
| t-test | **0,0325** | **0,0084** | 0,0101 | 0,7309 | 0,0608 | **0,0001** |

| Average intensity | Embryo_07 | Embryo_08 | Embryo_09 | Embryo_10 | Embryo_11 | Embryo_12 |
| --- | --- | --- | --- | --- | --- | --- |
| **PE cells** | 63,69 | 60,99 | 115,45 | 64,67 | 56,23 | 107,03 |
| St-Dev | 26,92 | 18,28 | 6,52 | 11,43 | 10,46 | 21,14 |
| **Pericardial cells** | 28,76 | 31,59 | 74,03 | 36,44 | 30,05 | 68,40 |
| St-Dev | 16,68 | 7,78 | 22,14 | 6,82 | 8,85 | 28,52 |
| t-test | **0,0055** | **0,0005** | **0,0001** | **0,0000** | **0,0002** | **0,0024** |

Movie 1. Multiple avcPE clusters in ift88 mutant.

Movie 2. Immotile and bent cilia protruding from the avcPE.

Movie 3. Absence of avcPE cluster in iguana mutant.

Movie 4. Yap/Wwtr1-Tead activity during avcPE development (51-60 hpf).

Movie 5. 3D projection of the PE (marked by anti-Wt1 antibody) of a control embryo at E 9.5.

Movie 6. 3D projection of the PE (marked by anti-Wt1 antibody) of a IFT20 KO embryo at E 9.5.

## Material and Methods

### Zebrafish (ZF) husbandry and embryo treatments

Animal experiments were approved by the Animal Experimentation Committee of the Institutional Review Board of the IGBMC. ZF lines used in the study were *Et(−26.5Hsa.WT1-1gata2:EGFP)^cn1^* transgenic line (referred to as *epi:GFP*) (46), *amotl2a ^fu46^* (33), *elipsa ^tp49d^* (10), *ift88 ^tz288/oval^* (53), *iguana ^ts294e^* (54), *yap1 ^fu48^* (33), *4xGTIIC:d2GFP* (61),*actb2:Mmu.Arl13b-GFP* (56) and *tcf21:NLS-EGFP* (87). All animals were incubated at 28.5°C for 24h before treatment with 1-phenyl-2-thiourea (PTU) (Sigma Aldrich) to prevent pigment formation.

Verteporfin (Sigma Aldrich) was diluted to 5 µM in fish tank water with 0.0033% PTU, in which larvae were incubated in darkness for 5h at 28.5°C.

### ZF Immunofluorescence

Embryos were fixed at the desired stages in 4% paraformaldehyde (PFA) overnight at 4°C. After washing in 0.1% PBS Tween 20, embryos were permeabilized in 0.5% PBS Triton X-100 for 20 min at room temperature (RT). Samples were washed and then blocked (3% albumin from bovine serum (BSA), 5% goat serum, 20 mM MgCl2, 0.3% Tween 20 in PBS) during 2h at RT. Primary antibodies were added in the blocking solution and incubated overnight at 4°C. Secondary antibodies were added in 0.1% PBS Tween20 after thorough washing and incubated overnight at 4°C. Embryos were washed and incubated with DAPI (Invitrogen), 1:1000, for 15 min at RT. After being thoroughly washed, samples were mounted for imaging on a Leica SP8 confocal with a dipping immersion objective (Leica HCX IRAPO L, 25X, N.A. 0.95). Z-stacks were taken every 10 µm. 3D images were reconstructed using IMARIS software (Bitplane Scientific Software). The ventral pericardium was digitally removed to provide a clearer view of the heart.

Antibodies used were as follows: anti-myosin heavy chain (MF20, DSHB) 1:20, anti-GFP (AVES) 1:500, anti-phospho-Smad 1/5 (Ser463/465) (Cell signaling) 1:50, anti-Yap1 (Lecaudey lab) 1:200. Secondary antibodies: goat anti-chicken Alexa Fluor 488 IgY (H+L) (In vitrogen), goat anti-mouse IgG Cy3 conjugate (H+L) ((Life technologies) and goat anti-rabbit Alexa Fluor 647 (ThermoFisher) were used at 1:500.

To test the effects of Verteporfin treatment, embryos were rinsed in fish tank water before being fixed and processed as described above.

### avcPE cell quantification

We performed whole-mount immunofluorescent staining on control and mutant embryos in the *epi:GFP* reporter line background and imaged the heart using a confocal microscope with a z-step of 10 µm. The different tissues were labeled using anti-myosin heavy chain (MHC) (myocardium) and anti-GFP (*epi:GFP*) antibodies, and DAPI dye to stain for nuclei. We then manually quantified the number of avcPE cells per z slice. We identified the avcPE clusters anatomically: avcPE clusters form in the dorsal pericardium, close to the atrio-ventricular canal (avc). PE cells were identified by their rounded morphology in conjunction with their expression of GFP (although some pericardial cells are also GFP-positive, they can be excluded due to their flat morphology). In order to count each cell only once in the z-stack, we only counted a cell when its nucleus was visible.

### Myocardial cell quantification

We performed whole-mount immunofluorescent staining on control and mutant embryos using anti-myosin heavy chain (MHC) (myocardium) antibody and DAPI dye (nuclei). We imaged the heart using a confocal microscope with a z-step of 10 µm. We then manually quantified the number of myocardial cells per z slice (nuclei surrounded by MHC signal).

### In situ hybridization (ISH)

ISH was performed in whole embryos according to (Thisse and Thisse, 2008) with minor modifications. Antisense mRNA probe used was against full coding sequence of *bmp4*.

### In vivo imaging

ZF embryos were staged, anaesthetised with 0.02% tricaine solution and mounted in 0.7% low melting-point agarose (Sigma Aldrich). Confocal imaging was performed on a Leica SP8 confocal microscope. Images were acquired bidirectionally with a low-magnification water immersion objective (Leica HCX IRAPO L, 25X, N.A. 0.95). For time lapse, z-stacks were acquired each 15 or 30 min, depending on the experiment. The optical plane was moved 15 µm between z-sections.

Bright field experiments were performed on a Leica DMIRBE inverted microscope using a Photron SA3 high speed CMOS camera (Photron, San Diego, CA) and water immersion objective (Leica 20X, NA 0.7). Image sequences were acquired at a frame rate of 150 frames per second.

### Mouse models

*Ift20^null/+^* (88) and *Ift88^null/+^* (89) mice were maintained on a C57BL/6JRj and B6D2 genetic background respectively. Animal procedures were approved by the ethical committee of the Institut Pasteur and the French Ministry of Research. E9.5 embryos were isolated in 200ng/ml cold heparin, incubated in cold 250mM KCl and fixed in 4% paraformaldehyde in PBS inside a rotative oven at 37°C overnight to remove excess of blood. Male and female samples were mixed.

### Whole mount immunofluorescence in the mouse

Embryos were fixed in paraformaldehyde. The cardiac region was dissected, permeabilised in 0.75% Triton. Aldehydes were quenched with 2.6mg/ml NH4Cl. Immunostaining was performed in 10% inactivated horse serum, 0,5% Triton with a primary antibody against Wt1 (Santa Cruz sc-192, 1:50), and with Alexa Fluor conjugated secondary antibodies (1:300) and counterstained with Hoechst (1:400). 80% glycerol was used to make the samples transparent.

### Mice PE volume analysis

Whole mount embryos stained with Wt1 antibody (Santa Cruz) were scanned on TCS SP8 DLS (Digital Light Sheet) Leica with a water immersion objective (HC APO L, 10X, 0.3). Z-stacks were taken every 2 µm. Both, coronal and sagittal views were acquired, if possible, for a more precise analysis. (**Supp. 2 F**) Using Matlab software, the contour of the PE was manually drawn (Wt1 signal) for each z plane and the area (A) was calculated. Volume (V) was estimated as: V = Σzn Ai x dz; i=1; zn=number of planes; dz= 2 µm 3D images were reconstructed using IMARIS software (Bitplane Scientific Software).

### Immunofluorescence on cryosections in the mouse

Embryos were embedded in 7% gelatin, 15% sucrose, frozen in cold isopentane and sectioned on a cryostat (10 µm). Immunostaining was performed on cryosections as described above, with permeabilisation in 0.5% Triton, and with an additional incubation in 0.2 mg/mL goat anti-mouse IgG Fab fragment to reduce non-specific reactivity of antibodies raised in the mouse. Primary antibodies against Tbx18 (Santa Cruz sc-17869, 1:100), Wt1 (Santa Cruz sc-192, 1:50), Yap1 (Santa Cruz sc-101199, 1:100), Amotl1 (Sigma HPA001196, 1:50) and p-Smad1/5/9 (Cell signaling 13820, 1:250) were used, with Alexa Fluor conjugated secondary antibodies (1:500) and Hoechst nuclear counterstaining (1:1000). Samples were imaged in DAKO mounting medium on a LSM700 (Zeiss) confocal microscope with a 40X/1.3 objective. Z-stacks were taken every 0.9 µm.

### Yap1− and Amotl1-positive cell quantifications

Nuclei positions on the slides were defined using IMARIS (Bitplane Scientific Software) Spots detection function. The results were manually corrected if needed. Nuclei positions were exported from Imaris and imported to Matlab. Each cell was assigned a unique index. The intensity of Tbx18 signal was evaluated in correspondence with the nuclei positions. Cells where the intensity was found higher than a threshold were considered positive. The threshold was established according to the background noise intensity. Results were manually corrected if needed and Tbx18-positive cells were automatically counted. Yap1 signal was visualized in fire LUT to facilitate perception of signal intensity. Nuclear Yap1-positive cells (higher signal in the nucleus than in the cytoplasm) were manually defined through index identification of cells and counted automatically. The same procedure was followed for Amotl1 signal. The outline of the outer PE region was manually drawn and areas of the two regions were calculated automatically. Matlab provided the total positive cell number for each signal and area, including signal co-localization (**Supp. 3 E**).

### Cell culture, siRNAs and transfection

Cells were cultured in appropriate conditions: MDCK (MEM Eagle - Earle’s BSS, 10 % FCS, AANE 0.1 mM, Sodium Pyruvate 1mM, Gentamicin 40 µg/ml), HeLa (DMEM 4.5 g/l glucose, 10% FCS, Penicillin 100 UI/ml, Streptomycin 100 µg/ml), HEK293 cells (DMEM 1g/L glucose, FCS 10%, Penicillin 100 UI/ml, Streptomycin 100 µg/ml) and DLD-1 (DMEM 4.5 g/l glucose, 10% FCS, Penicillin 100 UI/ml, Streptomycin 100 µg/ml)siRNA (Dharmacon) ON-Target plus - Control pool Non-targeting (D-001810-10-05) and SMART pool human IFT88 (L-012281-01) were used at 50 nM working concentration. Cells were transfected 16h after splitting using Opti-MEM medium and Oligofectamine reagent.

### Generation of DLD-1 IFT88-AID targeted cells

DLD-1 IFT88-AID cells were generated by adding an AID tag followed by a YFP tag at the 3′ end of the last exon on the IFT88 genomic locus. In detail, a clonal population of DLD-1 cells stably expressing TIR1-9xMyc protein was used for targeting (90). sgRNA targeting two regions adjacent to the 3′ end of IFT88 gene were introduced under the control of U6 transcription promoter into two separate vectors encoding for the expression of the Cas9 nickase (D10A) (91) (addgene 42335). A donor construct containing ≈ 600 bp recombination arms surrounding the 3′ end of IFT88 locus, in frame with a sequence encoding for an AID-YFP-Stop sequence, was generated. All three vectors were transfected into DLD-1 TIR1 cells using Xtreme Gene 9 DNA transfection reagent (Roche). Cells were sorted based on their YFP fluorescence and single clones were isolated. Homozygous targeted clones were identified by PCR. Targeting of IFT88 and degradation of IFT88-AID-YFP was confirmed by immunoblot following addition of Auxin (Sigma-Aldrich) at 500 µM in the culture medium for the indicated times.

### Immunofluorescence on cells

Cells were fixed in 100% MeOH for 6 min at −20°C (Ift88 and γ-tubulin antibodies), in 4% PFA for 7 min at RT (DLD-1 cells. Yap1 antibody) or in PFA 4% for 17 min at RT (MDCK and HeLa cells. Yap/Taz antibody). After washing in 0.1% PBS Tween20, cells were permeabilized in 0.5% PBS-NP40 and blocked in 5% BSA 1h at RT. Primary antibodies were added in the blocking solution and incubated overnight at 4°C. Secondary antibodies were added in 0.1% PBS Tween20 after washing and incubated for 2h at RT. Then cells were incubated with DAPI (Invitrogen), 1:1000, for 15 min at RT. After being thoroughly washed, samples were mounted for imaging on a Leica SP5 (siRNA experiments) or SP8 (DLD-1 experiments and experiments to assess subcellular localization) confocal microscope with an oil immersion objective (Leica HCX PL APO lambda blue, 63X, N.A. 1.4). Z-stacks were taken every 1 µm.

Antibodies used were as follows: anti-γ-tubulin (Santa Cruz) 1:500, anti-α-tubulin (Sigma, 1:2000), anti-IFT88 (Euromedex) 1:50, anti-Yap1 (4912) (Cell signaling) 1:50 (DLD-1 experiments), and anti-Yap/Taz (D24E4) (Cell signaling) 1:50 (MDCK and HeLa siRNA experiments). Secondary antibodies: goat anti-mouse IgG Cy3 conjugate (H+L) (Life technologies) and goat anti-rabbit Alexa Fluor 647 (In vitrogen) were used at 1:500.

### Nuclear Yap quantifications

Analyses were performed using Image J. Nuclei areas were selected manually using DAPI signal as reference on z-projection images (sum slices for DLD-1 experiment and maximum intensity projection for siRNA experiments). Yap average nuclear signal intensity was measured for the selected areas. In experiments in DLD-1 cells the values were measured for each individual nucleus, while in the case of siRNA experiments, all the nuclei in a slice were measured together. Values were normalized to their controls in order to merge data from different experiments.

### Nuclear/cytoplasmic Yap1 ratio

Analyses were performed using Image J. Nuclei areas were selected manually using DAPI signal as reference on z-projection images (sum slices). Cytoplasmic areas were selected using α-tubulin signal as reference. Yap average nuclear signal intensity was measured for the nuclear ROI. Yap average cytoplasmic signal intensity was measured after subtracting nuclear ROI from the cytoplasmic ROI.

### Lysates and immunoblotting

DLD-1 cell extracts were obtained after lysis with Laemmli sample buffer of an equal number of cells for each sample. Proteins were resolved by SDS-PAGE, transferred to nitrocellulose membranes and revealed by immunoblot using Western Lightning Plus-ECL kit (PerkinElmer).

### Immunoprecipitation (IP) assays

#### HEK293 cells

HEK293 cells (Q-BIOgene AES0503) were co-transfected with plasmids Flag-Amotl1 (92) and Ift20-GFP (9), or a Flag-control plasmid using Lipofectamine® 2000 Transfection Reagent (ThermoFisher SCIENTIFIC) and cultured for 48h. Proteins were extracted in a lysis buffer (10mMTris-Cl pH 7.5, 5mM EDTA, 150mM NaCl, 10% glycerol and 5% CHAPS) in the presence of protease inhibitors (cOmpleteTM Protease Inhibitor Cocktail, Roche). Immunoprecipitation of protein extracts was performed using a monoclonal anti-Flag antibody covalently attached to agarose (Anti-FLAG M2 Affinity gel, Sigma). Proteins were eluted in 2xNuPAGE LDS Sample Buffer (ThermoFisher). Proteins were separated on SDS– polyacrylamide gel electrophoresis and transferred to a nitrocellulose membrane. Proteins were detected with the primary antibodies against Flag (1:1000, Sigma F7425), GFP (1:1000, ThermoFisher CAB421), Ift20 (1:500, Proteintech, 13615-1-AP), Yap1 (1:1000, Cell Signaling 4912S) and Amotl1 (1:1000, Sigma HPA001196), followed by HRP-conjugated secondary antibodies (1:5000, Jackson ImmunoResearch) and the ECL detection reagent.

#### HeLa cells

We performed IPs using GFP-Trap (ChromoTek) agarose beads in two conditions: Control IP (YAP1-Myc (93), pEGFP-C1 and HA-Amotl1(23)) and IFT88 IP (YAP1-Myc, IFT88-GFP (94) and HA-Amotl1). HeLa cells (2x 10cm dish/ condition) were transfected with Lipofectamine 2000. Experiments were performed using the following setup: Cells were seeded at high density into 10cm dishes and transfected 16 hrs after seeding at 95% confluency. Twenty-four hours post-transfection, cells were seeded into 15cm dishes in order to achieve culture of isolated cells (10x 15cm dish/ condition). Proteins were extracted 60 hrs post-transfection in a lysis buffer (10mM TrisHCl, pH7.5; 150mM NaCl; 0.5mM EDTA; 0.5% NP-40; protease inhibitors Complete). GFP beads were washed once in lysis buffer and incubated with 16 mg of the protein lysate for 16 hrs at 4 ⁰C. Beads were washed four times in buffer without detergent and proteins were eluted by boiling for 10 mins. The input (1%) and IP were analyzed using immunoblot and the membranes were probed with anti-GFP (Abcam), anti-Yap/TAZ (D24E4, Cell Signaling) and anti-HA (Sigma Aldrich) antibodies.

#### Statistics

We applied D’Agostino & Pearson and Shapiro-Wilk normality tests to assess whether the samples fit a normal distribution and F test to compare variances. For normal distributed and homoscedastic samples, we used t-test or ANOVA. For non-parametric samples, we applied Man-Whitney or Kruskal-Wallis. The pertinent statistical analyses for each experiment were performed using GraphPad Prism 7 software. For the analysis of nuclear p-smad 1/5/9 and YAP1 signal in mice we used the non-parametric Chi-squared test of homogeneity to test whether the observed frequency of positive nuclei was equally distributed across the *wild type* and mutant embryos.

